# Multiomic Profiling Reveals Metabolic Alterations Mediating Aberrant Platelet Activity and Inflammation in Myeloproliferative Neoplasms

**DOI:** 10.1101/2022.12.08.519689

**Authors:** Fan He, Angelo B. A. Laranjeira, Tim Kong, Alice Liu, Katrina J. Ashworth, Nina M. Lasky, Daniel A. C. Fisher, Maggie J. Cox, Mary C. Fulbright, Lilian A. Heck, LaYow Yu, Stephen M. Sykes, Angelo D’Alessandro, Jorge Di Paola, Stephen T. Oh

**Affiliations:** Division of Hematology, Department of Medicine, School of Medicine, Washington University School of Medicine, St. Louis, MO, USA; Division of Hematology & Oncology, Department of Pediatrics, School of Medicine, Washington University School of Medicine, St. Louis, MO, USA; Department of Biochemistry and Molecular Genetics, University of Colorado-Anschutz Medical Campus, Aurora, CO, USA; Immunomonitoring Laboratory, Center for Human Immunology and Immunotherapy Programs, Washington University School of Medicine, St. Louis, MO, USA; Department of Pathology and Immunology, Washington University School of Medicine, St. Louis, MO, USA

**Keywords:** myeloproliferative neoplasm, platelet, oxidative phosphorylation, PI3K/AKT/mTOR, alpha-ketoglutarate

## Abstract

Platelets from patients with myeloproliferative neoplasms (MPNs) exhibit a hyperreactive phenotype. Here, we found elevated P-selectin exposure and platelet-leukocyte aggregates indicating activation of platelets from essential thrombocythemia (ET) patients. Single cell RNA-seq analysis of primary samples revealed significant enrichment of transcripts related to platelet activation, mTOR and oxidative phosphorylation (OXPHOS) in ET patient platelets. These observations were validated via proteomic profiling. Platelet metabolomics revealed distinct metabolic phenotypes consisting of elevated ATP generation, accompanied by increases in the levels of multiple intermediates of the tricarboxylic acid (TCA) cycle, but lower alpha-ketoglutarate (α-KG) in MPN patients. Inhibition of PI3K/AKT/mTOR signaling significantly reduced metabolic responses and hyperreactivity in MPN patient platelets, while α-KG supplementation markedly reduced oxygen consumption and ATP generation. *Ex vivo* incubation of platelets from both MPN patients and *Jak2 V617F* mice with α-KG significantly reduced platelet activation responses. Oral α-KG supplementation of *Jak2 V617F* mice decreased splenomegaly and reduced hematocrit, monocyte and platelet counts. Finally, α-KG incubation significantly decreased proinflammatory cytokine secretion from MPN CD14+ monocytes. Our results reveal a previously unrecognized metabolic disorder in conjunction with aberrant PI3K/AKT/mTOR signaling, contributing to platelet hyperreactivity in MPN patients.

## Introduction

Philadelphia-negative chronic myeloproliferative neoplasms (MPNs), including polycythemia vera (PV), essential thrombocythemia (ET), and myelofibrosis (MF), are clonal hematopoietic disorders characterized by overproduction of mature blood cells such as erythrocytes, granulocytes, and/or platelets (1). MPN patients have significantly elevated risks of thrombosis, with an estimated pooled prevalence of overall thrombosis among patients with MPN of 20% at diagnosis (2). Cytoreductive therapies and aspirin are commonly used for the treatment of MPN patients to reduce thrombosis risk (3).

MPN-associated thrombosis is considered a multifactorial event involving the complex interplay of blood and endothelial cells, the coagulation cascade, *JAK2* mutation allele burden, and chronic inflammation, all of which likely contribute to the prothrombotic phenotype (4). Increased thrombin generation, elevated levels of procoagulant microparticles and soluble P-selectin are seen in MPN patients and suggest hyperreactivity of platelets (5–7). Platelets from homozygous *JAK2 V617F* knock-in mice, which exhibit a PV-like phenotype, have reduced aggregation and increased bleeding time, whereas heterozygous *JAK2 V617F* mice, which exhibit an ET-like phenotype, showed increased platelet aggregation and reduced bleeding time (8, 9). Chronic inflammation is a hallmark of MPNs regardless of subtype (10). Aside from its classical functions in hemostasis, platelets also contribute to thromboinflammatory processes by directly interacting with leukocytes (11). Elevated platelet-leukocyte aggregates (PLA) has recently been found to be an independent risk factor for MPN-associated thrombosis, indicating that platelets could be a mediator between thrombosis and inflammation (12–14). While it is known that thrombotic risk in MPNs is not directly correlated with the platelet count, the factors mediating aberrant activation of platelets in MPNs have not been systematically evaluated.

Human platelets contain mitochondria, which supply approximately 40% of energy during the resting state. Oxidative phosphorylation (OXPHOS) increases significantly during platelet activation and secretion to meet the increased demand for energy (15). The dysregulation of platelet metabolism has been reported in several diseases, such as diabetes mellitus, sepsis and pulmonary hypertension (15). Previously, we reported that tumor necrosis factor (TNF) induces mitochondrial dysfunction in mice, which was associated with platelet hyperreactivity, particularly during aging (16). We also reported increased mitochondrial mass and platelet hyperreactivity in platelets from patients with MPNs, a disease associated with chronic inflammation and significantly high levels of TNF (10, 16–18). However, despite this clear association, the mechanisms driving metabolic dysregulation in MPN patient platelets remain incompletely understood.

Here, we apply multiomic approaches in conjunction with functional interrogation of platelets from MPN patients to delineate mechanisms of platelet hyperreactivity. We confirm that MPN patient platelets exhibit increased activation and uncover metabolic alterations, such as in oxidative phosphorylation and mTOR signaling activation as critical contributors to this platelet hyperreactivity. In addition, across MPN primary samples and mouse models, we demonstrate that alpha-ketoglutarate (α-KG) supplementation suppresses platelet activation and megakaryopoiesis, in part due to inhibition of cellular metabolism and mTOR pathway signaling, thereby identifying a promising therapeutic strategy for MPN patients.

## Results

### ET patients show significantly increased P-selectin levels and platelet-leukocyte aggregates (PLA) formation

First, platelet activation was measured in peripheral blood from 25 MPN (ET = 17, MF = 8) patients and healthy individuals (HI, n = 7, Table 1). There were no significant differences of age and sex between ET patients and healthy controls. Platelets from ET and MF patients had significantly higher basal P-selectin, a marker of α-granule activation and secretion, compared to healthy controls, with no significant differences observed after platelet agonist thrombin receptor activator peptide 6 (TRAP6) stimulation (Figure 1A & 1B). We also observed increased PLA formation in ET peripheral blood samples (Figure 1C & 1D), consistent with previous reports (19). Platelets from ET patients had higher ⍺IIbβ3 integrin activation following TRAP6 stimulation but there was no difference at baseline (Figure 1E). P-selectin exposure positively correlated with ⍺IIbβ3 integrin activation and PLA formation as expected (Figure 1F-H & Table 2).

**Figure 1.**
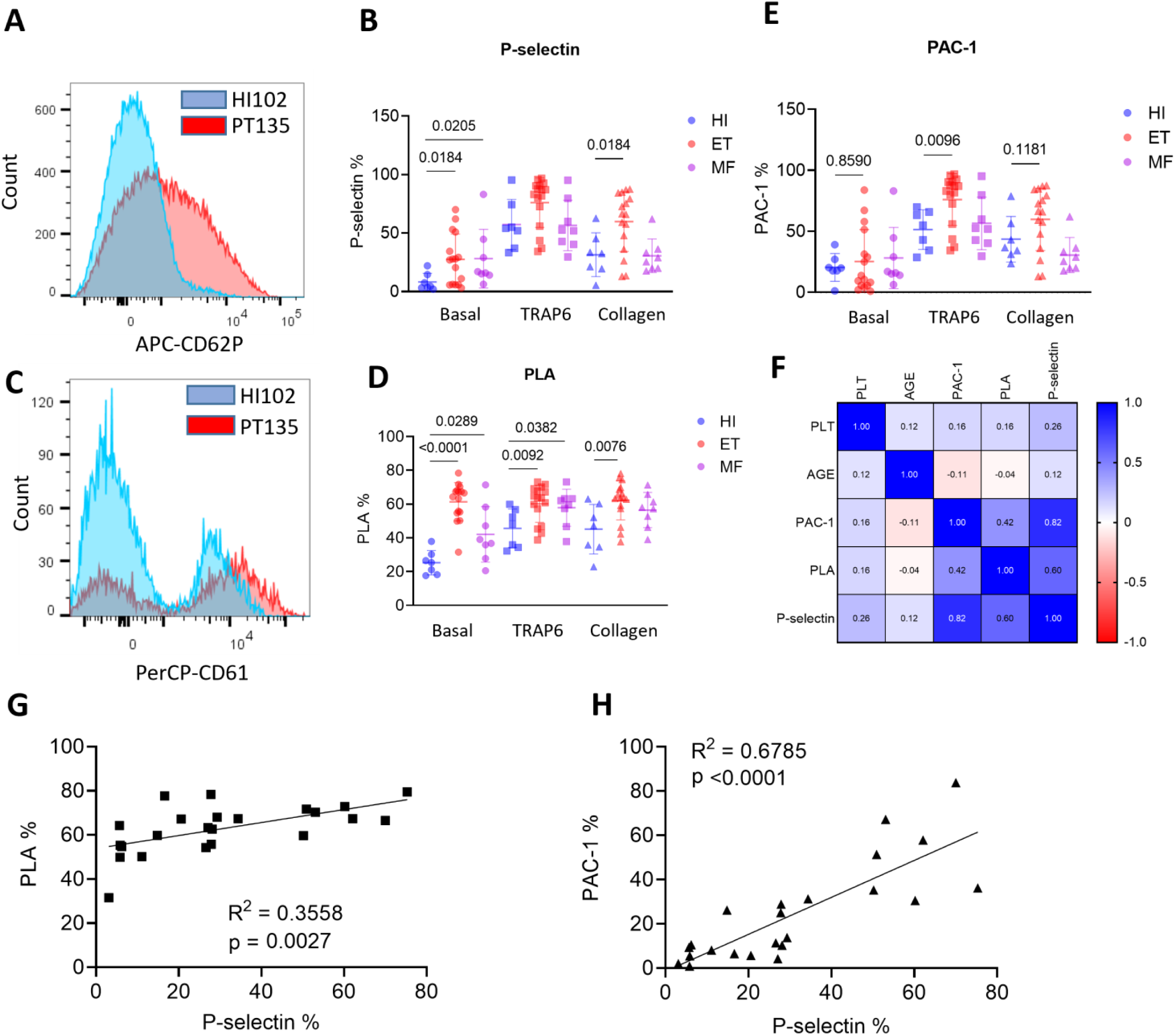
Platelets from ET patients show significantly increased P-selectin level and platelet-leukocyte aggregates (PLA) formation. A) Representative figure of exposure of P-selectin on the surface of platelets measured by flow cytometry. B) P-selectin expression on the surface of platelets at baseline and following 1 μM TRAP6 or 5 μg/ml collagen stimulation (HI = 7, ET = 17, MF = 8). Data are mean ± SD. Statistics were assessed by two-tailed Mann-Whitney U test. C) Representative figure of PLA ratio measured by flow cytometry. Data are presented as percents of aggregates from the respective leukocyte population. D) PLA measurements in whole blood at baseline and following 1 μM TRAP6 or 5 μg/ml collagen stimulation (HI = 7, ET = 17, MF = 8). Data are mean ± SD. Statistics were assessed by two-tailed Mann-Whitney U test. E) ⍺_IIb_β_3_ integrin expression (presented as the % positive staining of anti-PAC-1 antibody) at baseline and following 1 μM TRAP6 or 5 μg/ml collagen stimulation (HI = 7, ET = 17, MF = 8). Data are mean ± SD. Statistics were assessed by two-tailed Mann-Whitney U test. F) Pearson correlation coefficient among platelet markers and parameters. G) Simple linear regression between PLA % and P-selectin % in MPN patients. H) Simple linear regression between ⍺IIbβ3 integrin % and P-selectin % in MPN patients.

**Table 1.** Demographics of HIs and MPN patients for flow cytometry.

|  | JAK2 (n = 18) | CALR (n = 11) |
| --- | --- | --- |
| Age | 60.7 ± 13.7 | 63.5 ± 10.1 |
| Female % | 83.3% | 45.5% |
| PLT count | 508 ± 238 | 641 ± 327 |

**Table 2.** P value of Pearson’s correlations between platelet parameters from ET patients.

|  | PLT count | P-selectin % | PAC-1 % | Age | PLA % |
| --- | --- | --- | --- | --- | --- |
| PLT count |  | 0.228 | 0.469 | 0.601 | 0.474 |
| P-selectin % | 0.228 |  | 0 | 0.579 | 0.003 |
| PAC-1 % | 0.469 | 0 |  | 0.608 | 0.045 |
| Age | 0.601 | 0.579 | 0.608 |  | 0.861 |
| PLA % | 0.474 | 0.003 | 0.045 | 0.861 |  |

Platelets from patients carrying *JAK2* mutations exhibited stronger responses to TRAP6 stimulation than those with *CALR* mutations, as indicated by higher P-selectin exposure and ⍺IIbβ3 integrin activation; which is consistent with the lower risk of thrombosis reported in *CALR*-mutant patients compared to *JAK2* (Figure S1A & Table 3) (20). Platelet aggregometry showed decreased platelet responses in MPN patients when compared to HI, likely related to the effects of aspirin (Figure S1B & S1C). However, platelet aggregation responses were significantly higher in patients with the *JAK2* mutation when compared to those carrying *CALR* mutations (Figure S1D). Overall, our results show elevated P-selectin levels, increased ⍺IIbβ3 integrin activation and PLA formation indicating hyperreactivity of platelets in patients with MPNs.

**Table 3.** Demographics of the MPN patients for JAK2 and CALR comparison.

|  | HI (n = 7) | ET (n = 17) | MF (n = 8) |
| --- | --- | --- | --- |
| Age | 44.1 ± 14.7 | 55.6 ± 10.7 | 59.1 ± 7.4 |
| Female % | 42.9% | 76% | 75% |
| JAK2 mutation | NA | 12/17 | 3/8 |

### Platelets from ET patients show enrichment of genes involved in platelet activation, PI3K/AKT/mTOR signaling and oxidative phosphorylation (OXPHOS)

Based on our findings of increased platelet reactivity and PLA, and to better understand the transcriptional landscape of platelets and other blood cells from MPN patients, we performed single-cell RNA-seq (scRNA-seq) of peripheral blood samples from ET patients (n = 5) and age- and sex-matched HI (n = 3) (Figure 2A & Table 4). Cell types were assigned by their canonical transcripts, such as *CD14* for CD14+ monocytes, *CD8A* for CD8+ T cells and *PPBP* for platelets (Figure 2B & S2A). Cell clusters showed distinct transcriptional enrichments (Figure S2B). Subsequent cell composition analysis showed that CD4 T cells were the most abundant cell type in both ET and HI, accounting for 30%–55% of cells (Figure S2C). There were significantly increased percentages of platelets and a trend for higher monocyte counts in ET patients (Figure S2C). Platelets were further clustered into 10 distinctive groups using unsupervised clustering methods (Figure 2C & 2D). Platelets in cluster 6 exhibited the highest expression score for the platelet activation gene set and were all from ET patients, whereas cluster 0 with the lowest expression score was predominantly represented by HI platelets (Figure 2C & 2E). We then interrogated the overlap of the platelet activation gene set (composed of 261 genes) with differentially expressed genes in platelets from HI and ET. While none of the 261 genes were enriched in platelets from HI, 62 genes were upregulated in platelets from ET, including *SELP*, *PF4* and *GP1BA*, (Figure 2F & 2G). Consistently, unbiased GO and KEGG analyses showed enrichment of genes involved in platelet activation in ET patients (Figure S2D-S2F). Gene score enrichment analysis (GSEA) analysis confirmed increases in transcripts associated with platelet activation and IFN-γ pathways, and showed upregulation of genes involved in PI3K/AKT/mTOR signaling and OXPHOS in ET platelets (Figure S2G-S2K). We also performed transcription factor (TF) analysis using DoRothEA, a curated collection of TFs and their transcriptional targets (21). The enrichment of specific TFs in blood cells, such as *GATA1* in platelets and *ZEB2* in T cells, is consistent with previous studies (22, 23), which in turn support the accuracy of our cell identity assignments (Figure S2L & S2M). In addition, we observed upregulation of *GATA1* and *STAT1*, important regulators of megakaryopoiesis, in platelets from ET patients, which echoes elevated platelet count observed in ET patients (Figure S2N & S2O) (24).

**Figure 2.**
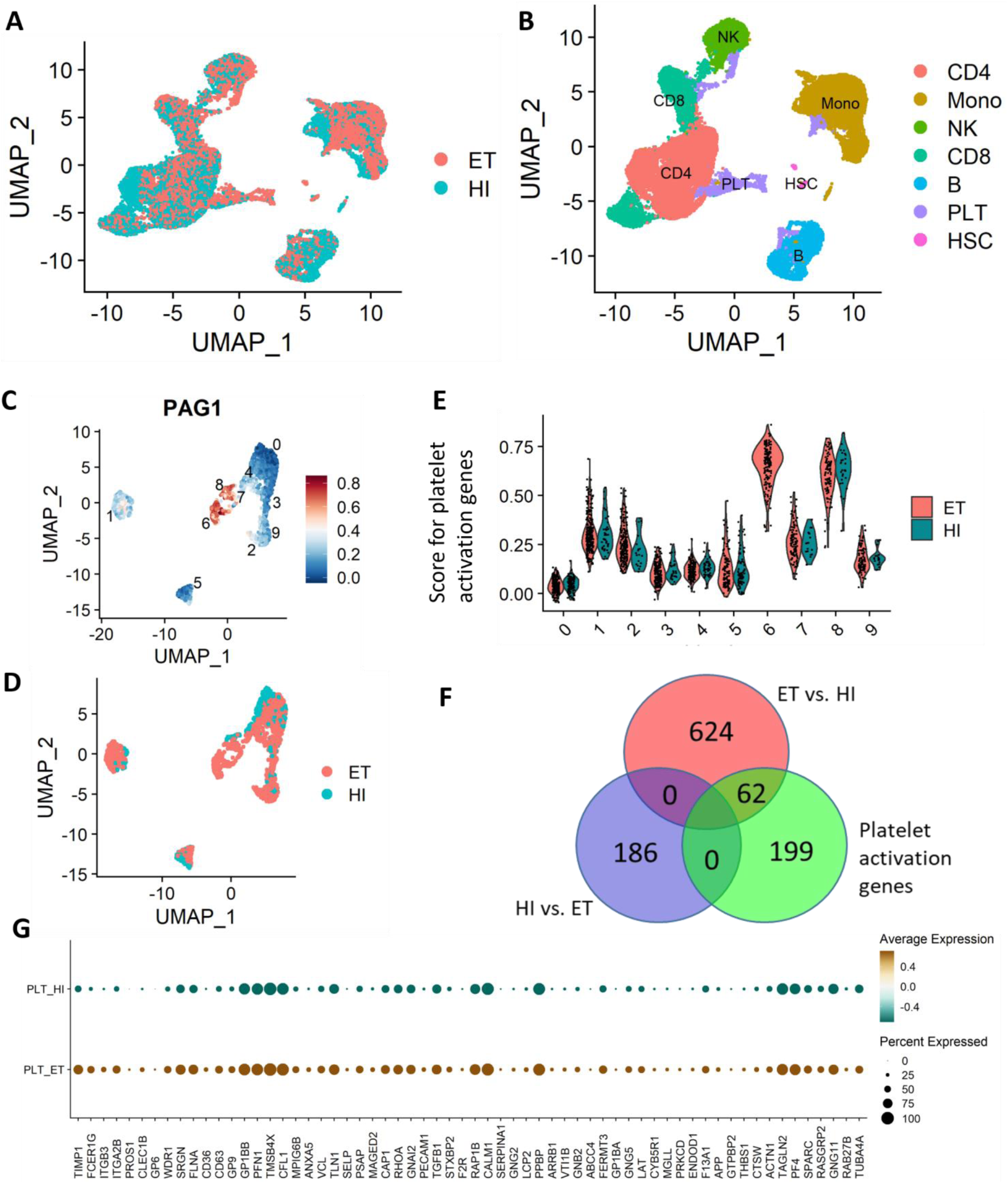
scRNA-seq revealed the activation of platelets and monocytes in peripheral blood from ET patients. A) UMAP plot of cells sequenced from HIs (n = 3) and ET patients (n = 5). B) UMAP plot of cells sequenced from HIs and ET patients with cell type annotations. C) UMAP plot showing platelets clustering with scores for “reactome platelet activation signaling and aggregation” gene set. D) UMAP plot showing platelets from HIs and ET patients. E) Violin plot of platelets clusters showing scores for “reactome platelet activation signaling and aggregation” gene set. F) Venn plot showing overlapped genes among differentially expressed genes in platelets from HIs, ET patients and genes in “reactome platelet activation signaling and aggregation” gene set. G) Dot plot of genes in “reactome platelet activation signaling and aggregation” gene set that overlaped with differentially expressed genes in platelets from HIs (0/186) and ET patients (62/686).

**Table 4.**
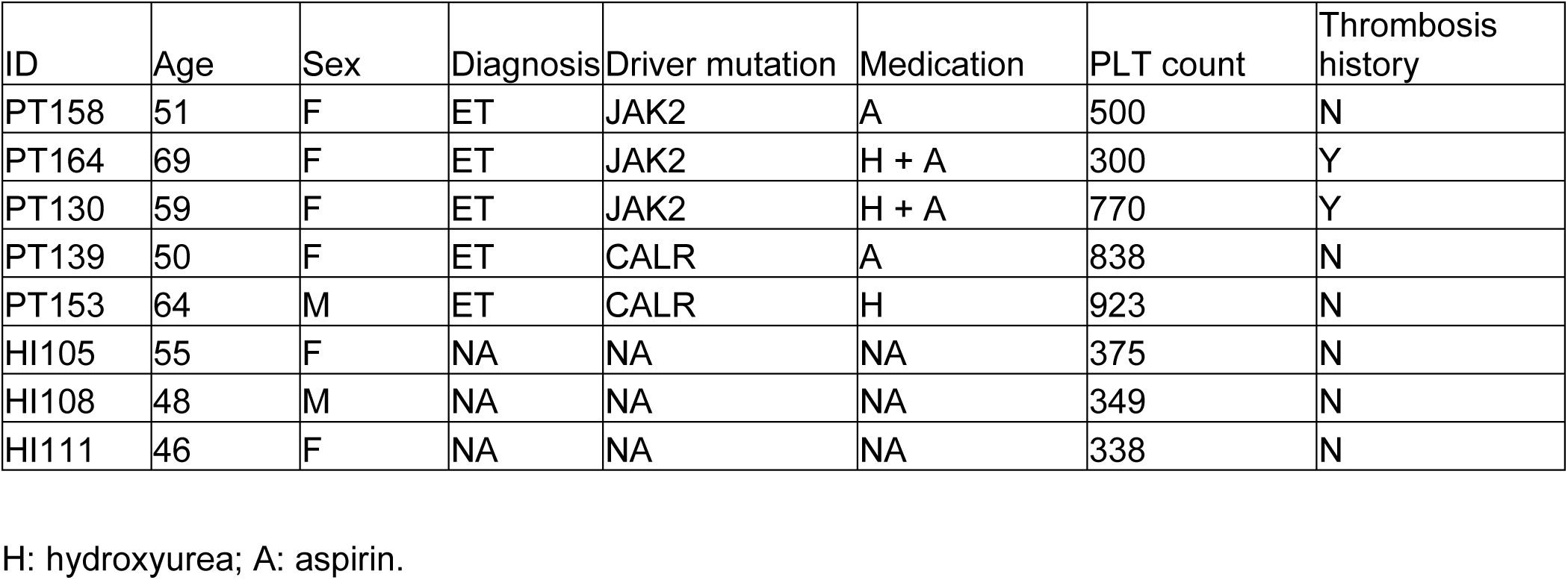
Demographics of HIs and MPN patients for scRNA-seq.

| ID | Age | Sex | Diagnosis | Driver mutation | Medication | PLT count | Thrombosis history |
| --- | --- | --- | --- | --- | --- | --- | --- |
| PT158 | 51 | F | ET | JAK2 | A | 500 | N |
| PT164 | 69 | F | ET | JAK2 | H + A | 300 | Y |
| PT130 | 59 | F | ET | JAK2 | H + A | 770 | Y |
| PT139 | 50 | F | ET | CALR | A | 838 | N |
| PT153 | 64 | M | ET | CALR | H | 923 | N |
| HI105 | 55 | F | NA | NA | NA | 375 | N |
| HI108 | 48 | M | NA | NA | NA | 349 | N |
| HI111 | 46 | F | NA | NA | NA | 338 | N |

H: hydroxyurea; A: aspirin.

To validate our scRNA-seq findings, we analyzed GSE2006 (25), a publicly available comparative microarray of platelets from ET and HI, and found enrichment of platelet activation and OXPHOS gene sets in platelets from ET patients (Figure S3A-S3D). Thus, these results not only support the hyperactivation of platelets in MPN patients, but also led to the hypothesis that platelet metabolic alterations contribute to the platelet hyperreactivity observed in MPNs.

Gene expression analyses of IFN-γ and inflammation response pathways revealed that monocytes exhibited the highest score of all blood cells analyzed (Figure S4A & S4C). Notably, monocytes from patients with *JAK2* V617F showed higher activities in IFN-γ and inflammation response pathways compared to patients with *CALR* mutations (Figure S4B & S4D). Enrichments of SPI1 and IRF TFs in monocytes from ET patients support our observations (Figure S4E). Further clustering showed that 2 distinctive monocyte clusters (8 and 9) exhibited the highest transcriptional levels of IFN-γ pathway genes (Figure S4F & S4G). Of note, these two clusters also showed robust enrichments of monocytes from ET patients and higher levels of *PPBP*, a platelet marker, suggesting a role of platelets in activating monocytes (Figure S4H & S4I). Thus, monocytes from ET patients display elevated inflammation and increased PLA formation, and may contribute to these responses.

### Metabolomics and proteomics analyses confirm elevated PI3K/AKT/mTOR signaling and mitochondrial activity in MPN platelets

To better understand metabolic alterations in MPN platelets, we performed ultra-high-performance liquid chromatography coupled with mass spectrometry (LC-MS) metabolomics on paired plasma and washed platelets from MPN patients along with age- and sex-matched HI (Figure 3A & Table 5, ET = 12, PV = 7, MF = 9, HI = 8). Platelet proteomics showed evident clustering of MPN samples in the PCA plot (Figure 3B). Metabolic signatures consistent with the enrichment of genes involved in mTOR signaling and OXPHOS pathways in platelets from MPN patients was also observed in metabolomics analyses (Figure 3C & 3D), consistent with our scRNA-seq data. We further performed multivariate analysis to reduce dimensionality for visualization, which revealed a clear distinction between HI and MPN platelets (Figure 3E). Differentially accumulated metabolites (DAMs) were defined as those exhibiting a log fold change > 0.5 and p < 0.05 between MPN and HI. In comparing MPN and HI, 24 of 181 (13.3%) were upregulated and 19 of 181 (10.5%) were downregulated; 6-Phosphogluconate, α-KG, succinate, and ATP were among top DAMs (Figure 3F & S5A). In contrast, the lack of differences in plasma metabolites between HI and MPN suggested that the metabolic alterations in platelets are intrinsic modifications instead of changes in microenvironment (Figure S5B & S5C). We further identified lower glucose in conjunction with higher pyruvate levels, indicating activation of glycolytic pathways in platelets from MPN patients compared to HI (Figure 3G). Increased Krebs cycle components succinate and malate paralleled with higher ATP levels, suggesting elevated mitochondrial activities in MPN patient platelets (Figure 3G). Measurement of mitochondrial protein and OXPHOS complexes by immunoblotting demonstrated significant increases of TOM-20, a mitochondrial marker, and complex V protein, the ATP synthase responsible for energy generation, in platelets from ET patients (Figure 3H & 3I).

**Figure 3.**
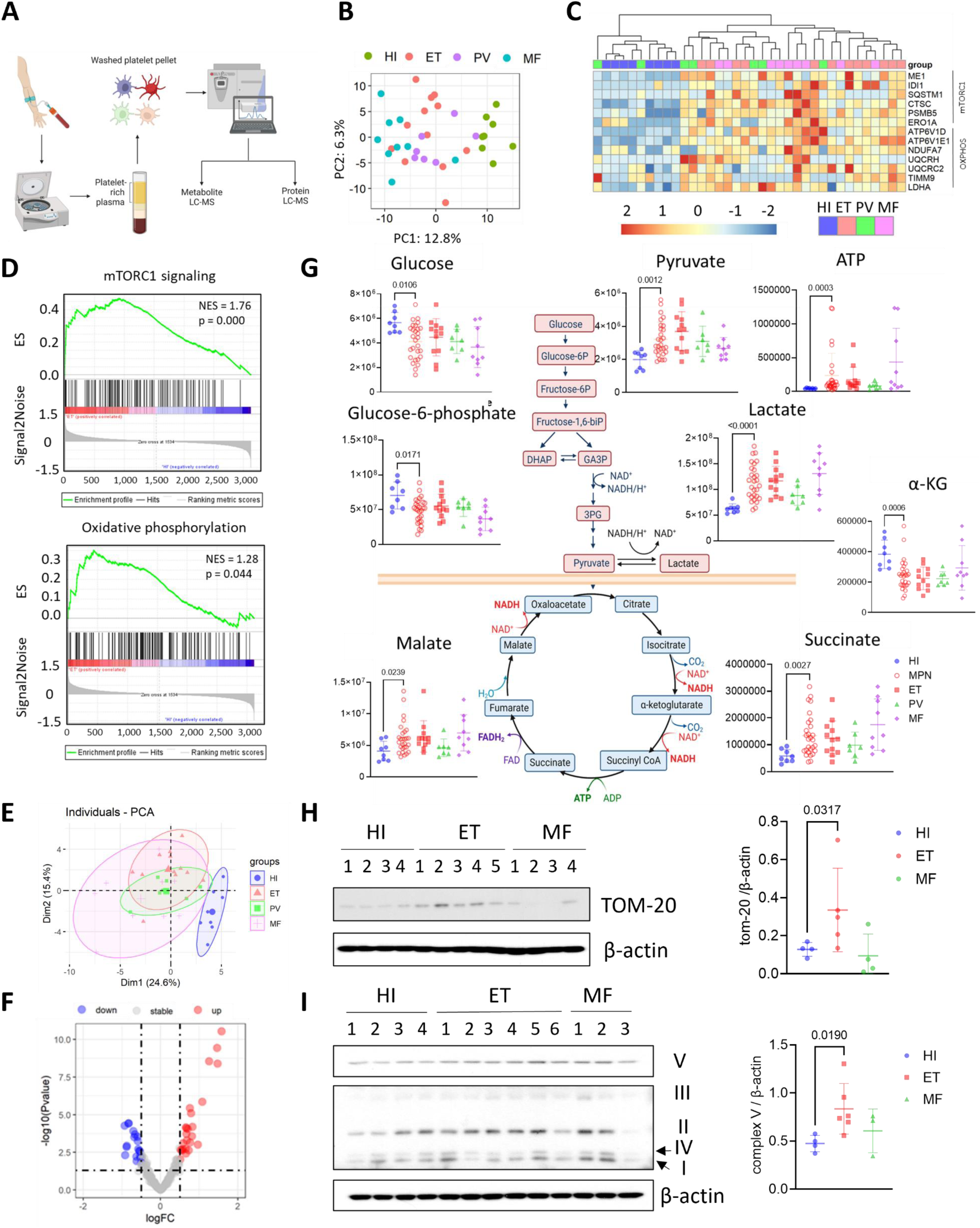
Metabolomics analyses showed distinct metabolic phenotypes of platelets from MPN patients. A) Diagram showing sample collection, processing and analysis. B) PCA scores plot of top 500 most variable proteins in protein LC-MS data of platelets from age- and sex-meatched HIs (n = 8) and MPN patients (ET = 12, PV = 7, MF = 9). C) Heatmap of selected proteins from “hallmark mTORC1 signaling” and “hallmark OXPHOS” gene sets. Columns are reorder based on the results of hierarchical clustering to identify sample correlations. D) GSEA enrichment plots for “hallmark mTORC1 signaling” and “hallmark OXPHOS” gene sets enriched in ET *vs.* HI. E) Principal component analysis (PCA) scores plot of metabolites LC-MS data of platelets from age- and sex-meatched HIs (n = 8) and MPN patients (ET = 12, PV = 7, MF = 9) displayed with 80% confidence region. F) Volcano plot of metabolite changes between HIs and MPN patients. Red dots denote significant (p < 0.05) and fold change (> 2^0.5) features, Blue dots denote significant (p < 0.05) and fold change (< - 2^0.5) features. G) The diagram showing steps of glycolysis and TCA cycle and scatter plots of peak areas (arbitrary units after normalization) for several key metabolites. Data are mean ± SD. Statistics were assessed by two-tailed Mann-Whitney U test. H) Western blot of washed platelets from HIs and MPN patients against TOM-20, a mitochondrial marker protein, and quantifications. Data are mean ± SD. Statistics were assessed by two-tailed Mann-Whitney U test. I) Western blot of washed platelets from HIs and MPN patients against human OXPHOS antibody cocktail and quantifications. Data are mean ± SD. Statistics were assessed by two-tailed Mann-Whitney U test.

**Table 5.** Demographics of HIs and MPN patients for platelet and plasma metabolomics and proteomics LC-MS.

|  | HI (n = 8) | ET (n = 12) | PV (n = 7) | MF (n = 9) |
| --- | --- | --- | --- | --- |
| Age | 51.5 ± 12.1 | 57.9 ± 11.2 | 54.9 ± 7.3 | 59.7 ± 5.6 |
| Female % | 37.5% | 58.3% | 42.9% | 88.9% |
| JAK2 mutation | NA | 8/12 | 7/7 | 4/9 |

### Inhibition of PI3K/AKT/mTOR signaling restrains MPN platelet hyperactivation

To investigate the role of PI3K/AKT/mTOR signaling in MPN platelet hyperactivation, we treated washed platelets from MPN patients with mTOR inhibitors. Omipalisib, a dual PI3K/mTOR inhibitor, abrogated all platelet activation and aggregation, whereas a milder, but still significant effect was observed by PI3K-sparring mTOR inhibitor sapanisertib (Figure 4A & 4B). In contrast, ruxolitinib, a JAK2 inhibitor approved for MPN treatment, had no effects on platelet activity. Further examination of platelet intracellular signaling pathways by immunoblotting revealed that mTOR inhibitors significantly reduced the phosphorylation of AKT and PLC-β induced by TRAP6 stimulation in a dose-dependent manner (Figure 4C & 4D). Incubation of omipalisib with platelets also completely blocked the energy demand boost induced by TRAP6 stimulation, which was not observed in sapanisertib or ruxolitinib (Figure 4E). Taken together, these results highlight key roles played by PI3K/AKT/mTOR signaling activation and elevated mitochondrial activity in MPN platelet hyperactivation.

**Figure 4.**
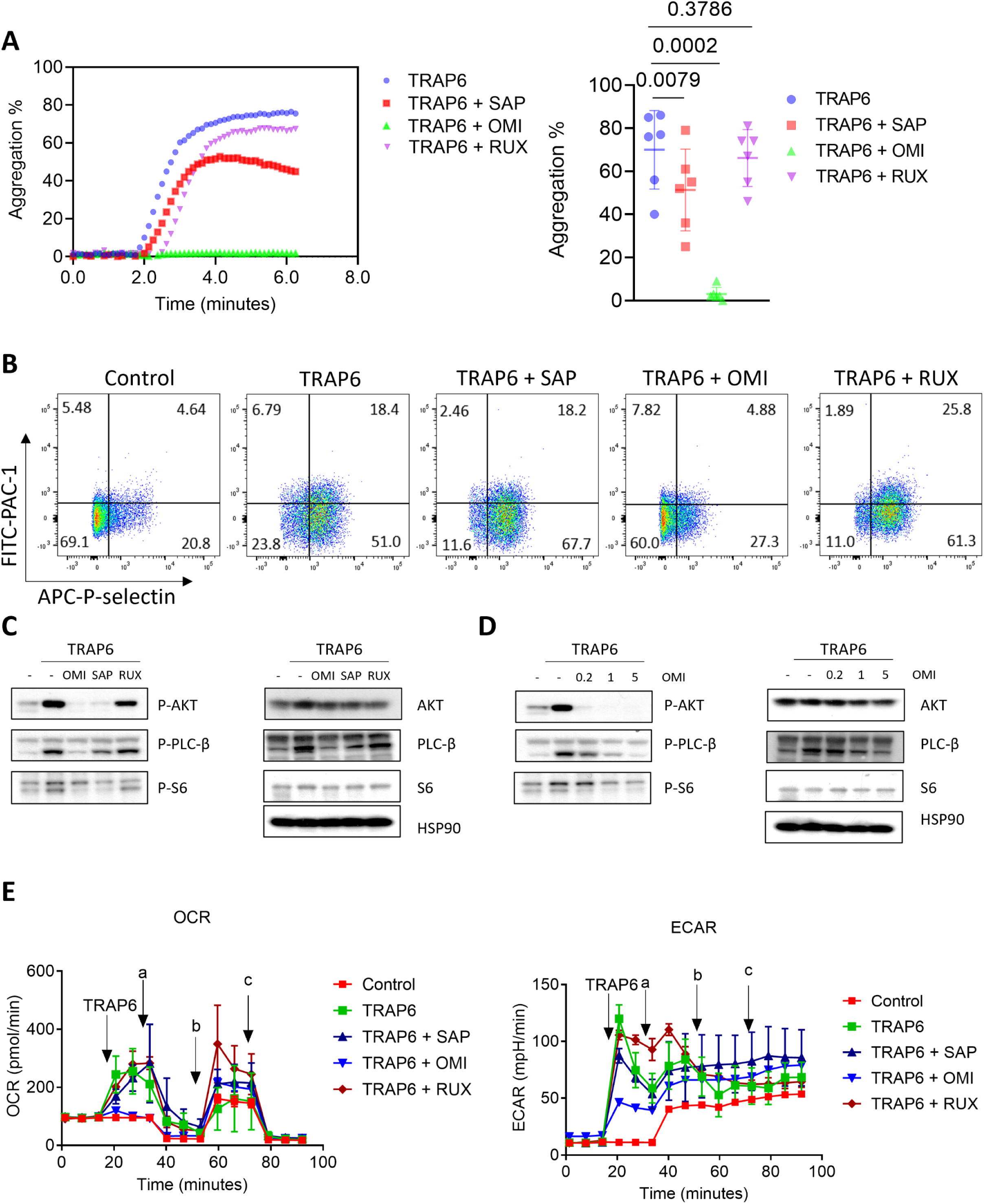
Effects of mTOR inhibitors and Ruxolitinib on platelet activities. A) Representative image and dot plot showing effects of mTOR inhibitors and Ruxolitinib on maximal aggregation intensity of washed platelets from ET patients. Washed platelets from ET patients were treated with Sapanisertib, Omipalisib or Ruxolitinib at 5 μM for 1 hour followed by platelet aggregatation analysis with 5 μM TRAP6 stimulation. Data is shown as mean ± SD. B) Representative images showing effects of mTOR inhibitors and Ruxolitinib on activation of washed platelets from ET patients. Washed platelets from ET patients were treated with Sapanisertib, Omipalisib or Ruxolitinib at 5 μM for 1 hour followed by flow cytometry analysis. C) Immunoblots showing changes of intracellular signaling pathways of platelets after mTOR inhibitors and Ruxolitinib treatments. Washed platelets from ET patients were treated with Sapanisertib, Omipalisib or Ruxolitinib at 5 μM for 1 hour followed by stimulation with TRAP6 peptides and immunoblot analysis. D) Immunoblots showing changes of intracellular signaling pathways of platelets after Omipalisib treatment. Washed platelets from ET patients were treated with Omipalisib at 0.2, 1 and 5 μM for 1 hour followed by stimulation with TRAP6 peptides and immunoblot analysis. E) Representative OCR and ECAR profiles of platelets from an ET patient showing the blockage of energy demand boost by mTOR inhibitors after TRAP6 stimulation. Washed platelets from the ET patient were treated with Sapanisertib, Omipalisib or Ruxolitinib at 5 μM for 1 hour followed by seahorse analysis (a: Oligomycin A; b: FCCP; c: Rotenone/antimycin A). 20 μM TRAP6 were injected on-plate to stimulate platelet energy demand.

### MPN platelets display bioenergetic alterations which can be reverted by α-KG supplementation

To interrogate bioenergetic alterations in platelets from MPN patients, platelets were isolated for Seahorse extracellular flux analysis. MPN platelets showed increased rates of basal respiration and ATP generation after correction for non-mitochondrial oxygen consumption rate (OCR); which suggested elevated physiological mitochondrial respiration (Figure 5A & 5B). Similar changes were observed in platelets from *Jak2 V617F* mice (Figure S6A & S6B). The reserve capacity, calculated as the difference between maximal and basal OCR, was also significantly greater in platelets from MPN patients (Figure 5B). Notably, *ex vivo* stimulation of platelets from MPN patients with TRAP6 showed greater OCR responses than in platelets from HI, indicating a larger reserve capacity in the setting of increased energy demand (Figure 5C). We further identified positive correlations of basal respiration with P-selectin exposure and maximum respiration, suggesting the importance of mitochondrial respiration and energy supply in platelet hyperactivation (Figure 5D). Altogether, our results demonstrate elevated mitochondrial respiration and coupling of ATP generation in MPN platelets.

**Figure 5.**
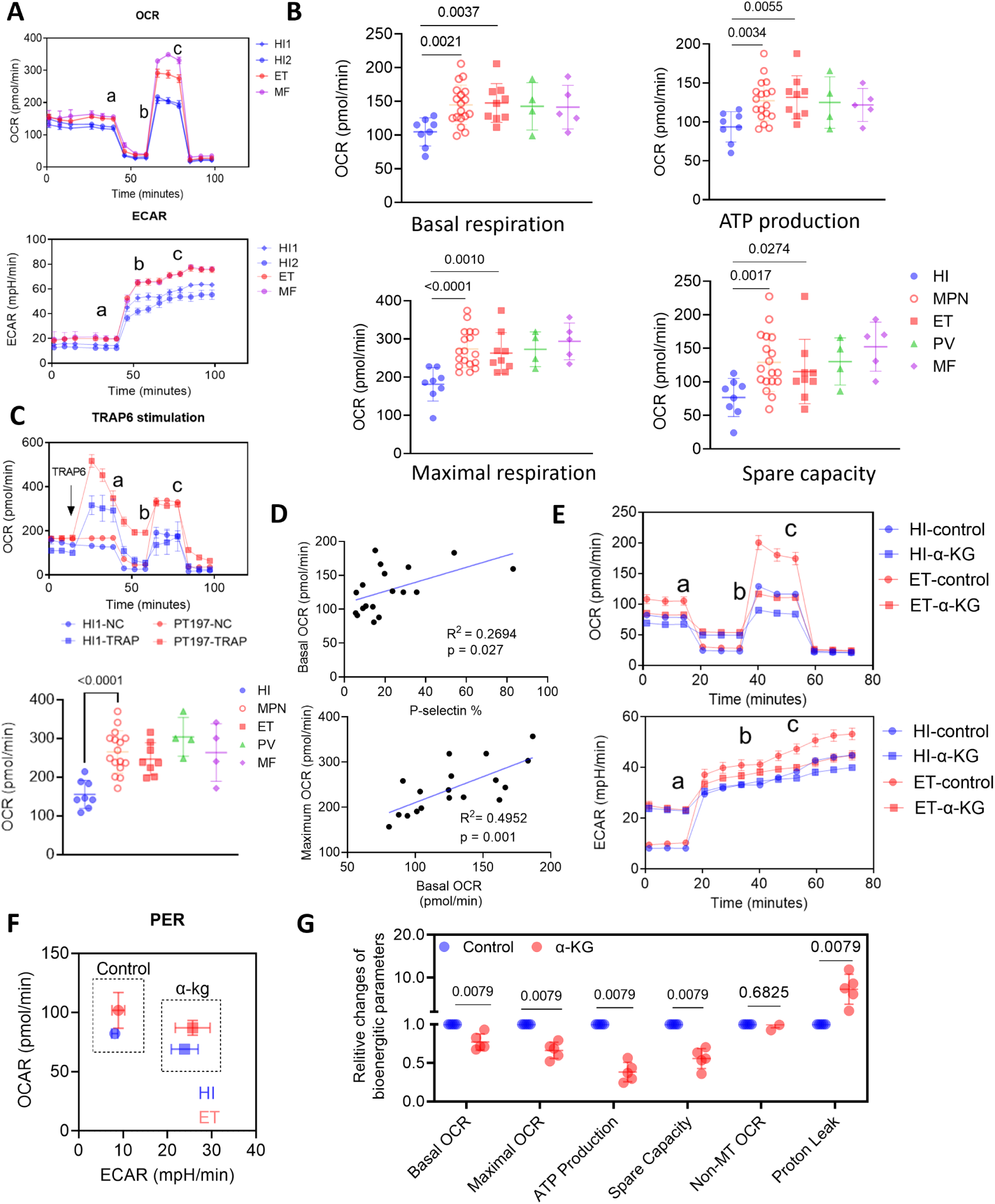
Platelets from MPN patients displayed bioenergetic alterations, which can be reverted by α-KG supplementation. A) Representative OCR and ECAR profiles of platelets from 2 HIs, a ET patient and a MF patient (a: Oligomycin A; b: FCCP; c: Rotenone/antimycin A). B) Quantification of basal OCR, ATP production, maximal OCR and spare capacity profiles of washed platelets in HIs (n = 8) and MPN patients (n = 28, ET = 12, PV = 7, MF = 9). Data are mean ± SD. Statistics were assessed by two-tailed Mann-Whitney U test. C) Representative OCR profiles of platelets from a HI and a ET patient showing the energy demand boost after TRAP6 stimulation and quantification of post-TRAP6 stimulation OCR profiles (a: Oligomycin A; b: FCCP; c: Rotenone/antimycin A). Data are mean ± SD. Statistics were assessed by two-tailed Mann-Whitney U test. D) Corelation analysis of bioenergetic parameters and platelet functional parameters. E) Representative OCR and ECAR profiles of platelets from a HI and a ET patient with the pre-incubation of 500 μM Octyl-α-KG or DMSO for 1 hour (a: Oligomycin A; b: FCCP; c: Rotenone/antimycin A). F) Platelet OCR/ECAR ratio from a HI and a ET patient with the pre-incubation of 500 μM Octyl-α-KG or DMSO control. G) Quantification of individual components of the platelet OCR profile in MPN patients (n = 5). Data were normalized to DMSO group set as 1. Data are mean ± SD. Statistics were assessed by two-tailed Mann-Whitney U test.

α-KG is a key intermediate of the TCA cycle, which is produced from isocitrate by oxidative decarboxylation or from glutamate by oxidative deamination (26). Recent studies found that α-KG supplementation inhibits ATP generation via direct suppression of ATP synthase and plays a critical role in alleviating aging and inflammation (27, 28). A recent report also showed that α-KG inhibits thrombosis and inflammation via suppression of AKT phosphorylation (29). Based on our findings of activation of OXPHOS and mTOR signaling along with decreased α-KG levels in MPN patient platelets, we tested whether α-KG supplementation could revert platelet metabolic alterations in the context of MPN. Indeed, α-KG incubation significantly reduced basal respiration, ATP generation and reserve capacity in MPN platelets (Figure 5E-5G). α-KG inhibited proton pump in ATP synthase as expected, reflected by increased proton leak as well as increased extracellular acidification rate (ECAR) (Figure 5E-5G). Thus, α-KG supplementation modified the metabolic phenotype of platelets from MPN patients by inhibiting mitochondrial activity. A similar inhibition of mitochondrial respiration by α-KG was observed in UKE-1, a *JAK2*-mutant ET transformed acute leukemia cell line (Figure S6C &S6D).

### α-KG suppresses PI3K/AKT/mTOR signaling and platelet activation through ATP synthase inhibition

After demonstrating the capacity of α-KG to inhibit platelet metabolism, we further evaluated the influence of α-KG on platelet activation. *Ex vivo* incubation of α-KG significantly reduced P-selectin expression and ⍺IIbβ3 integrin activation following TRAP6 treatment of washed platelets from MPN patients (Figure 6A). A similar inhibitory effect of α-KG was observed in platelets from *Jak2 V617F* mice (Figure S6E). Since αIIbβ3 downstream signaling (outside-in signaling) plays a critical role in platelet spreading and aggregation, we performed static platelet spreading and adhesion assays on platelets from *Jak2 V617F* mice. *Ex vivo* incubation with α-KG significantly decreased the number and area of platelets adhering to fibrinogen-coated surfaces; α-KG also inhibited platelet spreading induced by a PAR-4 agonist peptide (Figure 6B). Consistently, α-KG induced a dose-dependent inhibition of platelet aggregation (Figure 6C).

**Figure 6.**
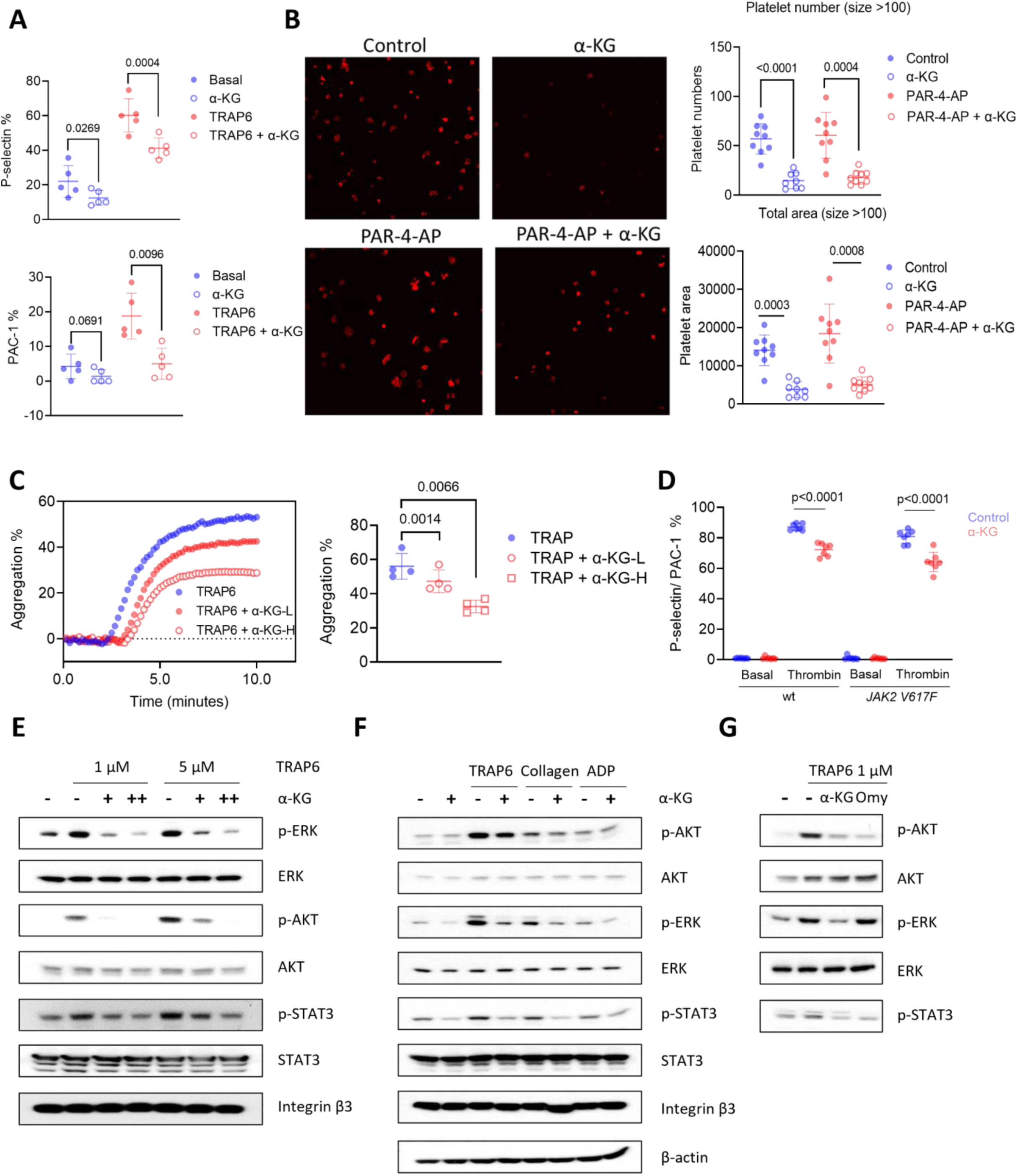
α-KG inhibited platelet activation through the suppression of ATP synthase. A) Platelet P-selectin and ⍺IIbβ3 integrin expression changes after incubation with 200 μM Octyl-α-KG for one hour. Data are mean ± SD. Statistics were assessed by two-tailed paired t test. B) Platelet adhesion and spreading assay with α-KG. Washed platelets from *VavCre*/*Jak2 V617F* knock-in mice were incubated with Octyl-α-KG at 37°C for one hour and/or PAR-4 agonist peptide for 5 minutes followed by placing on fibrinogen-coated coverslips for 45 minutes at 37°C. Number and area of attached platelets on the coverslips were quantified with CellProfiler software (n=9). Data are mean ± SD. Statistics were assessed by two-tailed Mann-Whitney U test. C) Platelets aggregation assay with α-KG treatment. Washed paltelets from peripheral blood of ET patients were incubated with 100 μM (L) or 200 μM (H) Octyl-α-KG for one hour and stimulated with 5 μM TRAP6 followed by platelet aggregation tests. Maximal aggregation intensity was quantified as mean ± SD. Statistics were assessed by two-tailed paired t test. D) Platelet P-selectin and ⍺IIbβ3 integrin expression changes in α-KG supplemented mice. Age- and sex-matched wild type and *VavCre/Jak2 V617F* knock-in mice were supplemented with regular water (n= 7) or 2% α-KG (n = 7) for one week. Platelets were stimulated with thrombin *ex vivo* or not followed by flow cytometry analysis. Data is shown as the ratio (mean ± SD) of P-selectin/⍺IIbβ3 integrin dual positive platelets. Statistics were assessed by two-tailed Student’s t test. E) Immunoblots of washed platelets after α-KG treatment. Washed platelets from ET patients were incubated with 250 μM or 500 μM Octyl-α-KG for one hour followed by stimulation with TRAP6 peptides. F) Immunoblots of washed platelets after α-KG treatment with different stimulant. Washed platelets from ET patients were incubated with 250 μM Octyl-α-KG followed by stimulation with 5 μM TRAP6 peptides or 5 μg/ml collagen or 20 μM ADP. G) Immunoblots of washed platelets after α-KG or Oligomycin (Omy) treatment. Washed platelets from ET patients were incubated with 250 μM Octyl-α-KG or 1 μM Omy followed by stimulation with TRAP6 peptides.

Supplementation with 2% α-KG in the drinking water inhibited P-selectin exposure and ⍺IIbβ3 integrin activation on platelets in both wild type and *Jak2 V617F* knock-in mice following thrombin stimulation (Figure 6D). Mechanistically, previous reports have shown that phosphorylation of STAT3, AKT and ERK play essential roles in platelet activation (30–32). Here, we found that α-KG inhibited p-STAT3, p-AKT and p-ERK1/2 in MPN patient platelets in a dose-dependent manner following TRAP6 stimulation, without affecting the total amount of these proteins (Figure 6E). Moreover, α-KG downregulated p-STAT3, p-AKT and p-ERK1/2 when platelets were activated with other agonists including collagen and ADP (Figure 6F). Oligomycin, a widely used inhibitor of ATP synthase, showed similar downregulation of p-AKT and p-STAT3, suggesting a role for ATP synthase in mediating platelet activation (Figure 6G) (33). Finally, addition of α-KG inhibited the proton pump of complex V in conjunction with decreased intracellular ATP levels but increased mitochondrial membrane potential in the megakaryocytic MEG-01 cells as well *as JAK2*-mutant UKE-1 cells (Figure S6F-S6I). Collectively, our data demonstrate that α-KG supplementation inhibits MPN platelet hyperactivation, in part due to inhibition of ATP synthase.

### α-KG exerts therapeutic effect on MPN and inhibits megakaryopoiesis

After demonstrating the effects of α-KG on reducing platelet hyperreactivity *ex vivo*, we next sought to investigate the therapeutic impact of α-KG in MPNs *in vivo*. Mice engrafted with *Jak2 V617F* c-Kit+ cells were supplemented with 1% α-KG in the drinking water vs. regular water for 6 weeks (Figure 7A). As expected, mice transplanted with *Jak2 V617F* cells developed characteristic MPN features including splenomegaly, increased white blood cell (WBC), red blood cell (RBC), and platelet counts (Figure 7B & 7C). α-KG supplementation markedly reduced splenomegaly, platelet, RBC and monocyte counts without affecting body weight or lymphocyte count (Figure 7B, 7C & S7A). Analysis of hematopoietic stem/progenitor cell (HSPC) subsets revealed a trend of decreased multipotent progenitor 2 (MPP2) cells in bone marrow from α-KG-supplemented mice, consistent with suppression of megakaryopoiesis and erythropoiesis (Figure S7B). Bone marrow histopathological analysis confirmed decreased megakaryocytes in α-KG supplemented mice (Figure 7D). Importantly, α-KG supplementation to wild type mice did not affect WBC or myeloid differentiation (Figure S7C).

**Figure 7.**
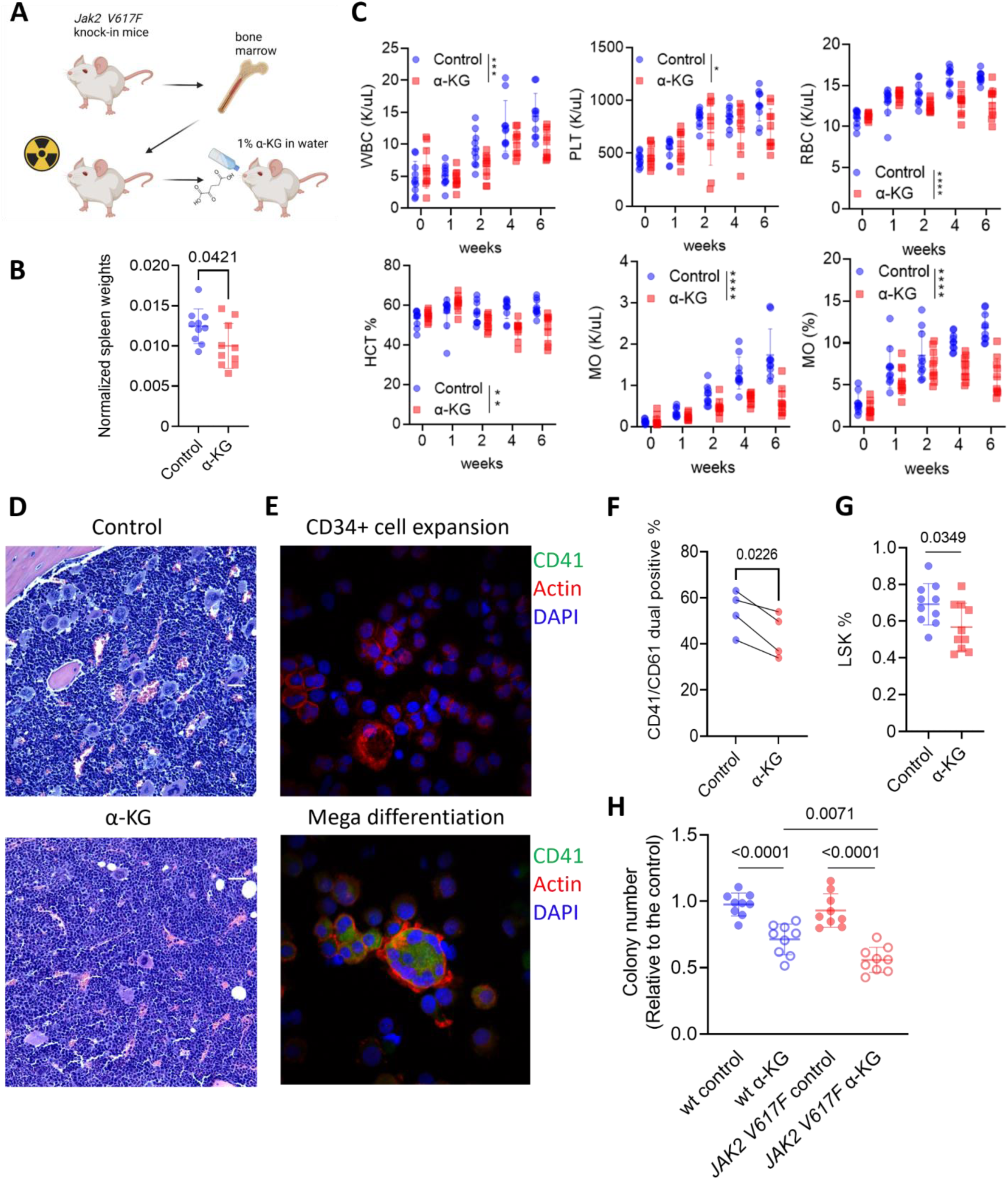
α-KG exerts therapeutic impacts on MPN and inhibits megakaryopoiesis. A) Schematic of the *JAK2 V617F* knock-in mice. cKIT+ cells from CD45.2 C57BL/6J mice were isolated and transplanted into irradiated CD45.1 C57BL/6J mice. Mice were supplemented with regular water (control, n = 10) or 1% α-KG in drinking water (n = 10) daily for 6 weeks. B) Spleen weight of transplanted mice normalized to body weight measured at the end of treatments. Data are mean ± SD. Statistics were assessed by two-tailed Student’s t test. C) WBC, platelet count, RBC, HCT, monocyte count and ratio of transplanted mice treated with regular water or α-KG across multiple timepoints. Data are mean ± SD. Statistics were assessed by two-way ANOVA with Dunnett’s multiple comparisons test (*P < 0.05; **P < 0.01; ***P < 0.001; ****P < 0.0001). D) Representative images of HE staining of femur bones from mice treated with regular water or α-KG. E) Representative images of immunofluorescence staining of expanded CD34+ cells and differentiated megakaryocytes from the same individual. F) Flow cytometry of CD41 and CD61 exposures on *in vitro* megakaryocytes differentiation with α-KG. Sorted CD34+ hematopoietic stem and progenitor cells were cultured for megakaryocytes differentiation with 250 μM Octyl-α-KG or DMSO control. CD41/CD61-dual positive cells were determined by flow cytometry. Data are mean ± SD. Statistics were assessed by two-tailed paired t test. G) Percentage of LSK cells of transplanted mice treated with DMSO control or α-KG at the end of treatments. Data are mean ± SD. Statistics were assessed by two-tailed Student’s t test. H) Colony-forming unit (CFU) assays of mouse LSK cells with DMSO control or α-KG. Colony numbers were counted after 14 days. Data are mean ± SD. Statistics were assessed by two-tailed Student’s t test.

To further characterize the effects of α-KG on megakaryopoiesis, sorted human CD34+ cells were differentiated into megakaryocytes *ex vivo* with the addition of cytokines (Figure 7E). Decrease CD41+/CD61+ dual positive megakaryocytes were observed in the presence of α-KG, consistent with inhibition of megakaryopoiesis in α-KG supplemented-mice (Figure 7F). Moreover, α-KG inhibited megakaryocyte maturation of MEG-01 cells *in vitro* (Figure S7D & S7E). Notably, α-KG treatment also reduced Lin−Sca-1+c-Kit+ (LSK) cells in mice, suggesting a role of α-KG in HSPC regulation (Figure 7G). Finally, α-KG led to a reduction in myeloid colony formation from both wild type and *Jak2 V617F* c-Kit+ cells, but to a larger degree in *Jak2 V617F* cells (Figure 7H).

RNA sequencing analysis revealed that bone marrow cells of α-KG-supplemented mice clustered separately from control mice, suggesting differential gene expression induced by α-KG supplementation (Figure S8A). GSEA analysis showed inhibition of OXPHOS, mTORC1, and myeloid differentiation pathways in α-KG mice (Figure S8B-S8E). Expression of *Gata1* and *Epor*, critical mediators of platelet and RBC differentiation were decreased in α-KG-supplemented mice (Figure S8F). Downregulation of *Cdk1* and *Cdkn3* in α-KG-supplemented mice suggested cell cycle arrest (Figure S8F). These findings collectively indicate that α-KG suppresses MPN disease features and megakaryopoiesis in *Jak2 V617F* mice.

### α-KG inhibits monocyte activation and hyperinflammation

While MPN is initiated by the acquisition of mutations in HSPCs, its progression is driven, at least in part, by inflammation (10). We found significant elevation of pro-inflammatory cytokines in plasma from MPN patients, including TNF, IFN-γ, IP-10 and IL-6, compared to HI (Figure 8A & Table 6). Previously, our group showed that monocytes play a critical role in hyperinflammation that characterizes MPNs (17, 18), many of which correlate with prognosis (34). As such, since monocyte counts decreased as early as two weeks in α-KG-supplemented mice (Figure 7C), we sought to investigate if α-KG impacted monocyte activation and inflammation.

**Figure 8.**
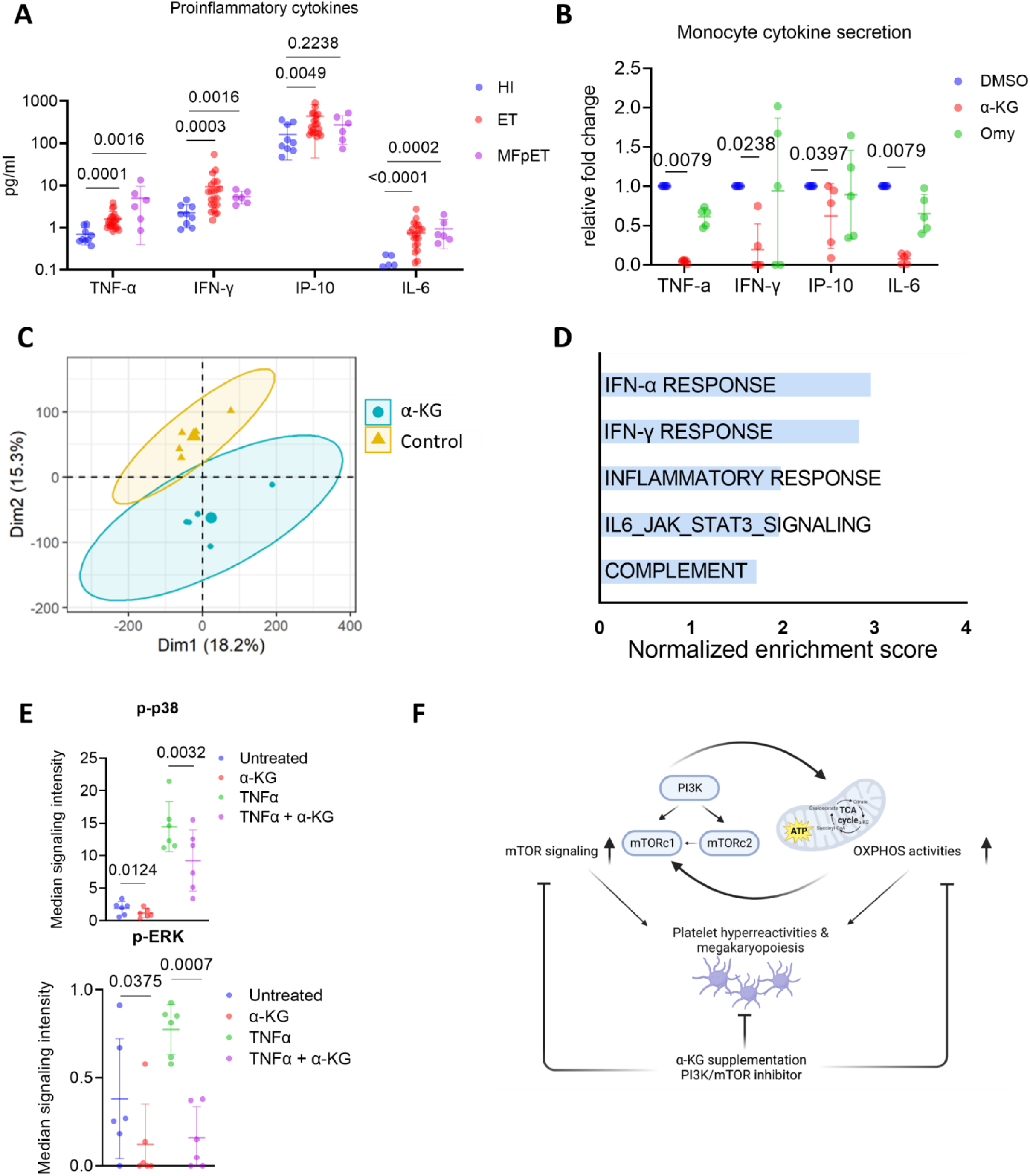
α-KG inhibited monocytes activation and hyperinflammation. A) Plasma cytokine levels in HIs and MPN patients. Plasma from HIs (n = 9) and MPN patients (n = 28) were collected for the determination of 30 biomarkers using V-PLEX Human Cytokine 30-Plex Kit from Meso Scale Discovery. Data are mean ± SD. Statistics were assessed by two-tailed Mann-Whitney U test. B) Monocyte cytokine secretion changes with Octyl-α-KG or Omy treatment. 0.5M sorted CD14+ monocytes from MPN patients (n = 5) were incubated with Octyl-α-KG or Omy for 8 hours and the supernants were collected for cytokine determination using V-PLEX Human Cytokine 30-Plex Kit. Data are mean ± SD. Statistics were assessed by two-tailed Mann-Whitney U test. C) The PCA scores plot of RNA-seq of sorted CD14+ monocytes after the incubation with Octyl-α-KG or DMSO control. D) Barplot of top 5 hallmark pathways with the highest enrichment scores in GSEA analysis. E) Dot plots of altered intracellular pathways of monocytes in peripheral blood of MPN patients by mass cytometry. Whole blood from MPN patients (n = 6) were incubated with Octyl-α-KG or DMSO for 1 hour followed by the stimulation of TNFα. Samples were processed for the determination of intracellular pathway activities by CyTOF. Data are mean ± SD. Statistics were assessed by two-tailed paired t test. F) Proposed model showing the roles of PI3K/AKT/mTOR signaling and metabolic alterations in platelets from MPNs. A positive feedback loop involving PI3K/AKT/mTOR signaling and metabolic alterations promotes platelet hyperreactivities and megakaryopoiesis in MPN. The supplementation of α-KG, which disrupts the feedback loop, shows therapeutic effects against platelet hyperreactivity, megakaryopoiesis and chronic inflammation in MPN.

**Table 6.** Demographics of HIs and MPN patients for plasma cytokine screening.

|  | HI (n = 9) | ET-JAK2 (n = 11) | ET-CALR (n = 11) | MFpET (n = 6) |
| --- | --- | --- | --- | --- |
| Age | 54.7 ± 20.1 | 57.5 ± 20.0 | 58.5 ± 17.6 | 64.5 ± 7.8 |
| Female % | 44.4% | 72.7% | 45.5% | 66.7% |

*Ex vivo* incubation of α-KG with sorted CD14+ human monocytes significantly decreased the secretion of pro-inflammatory cytokines (Figure 8B). Bulk RNA-seq results demonstrated that α-KG downregulated genes involved in interferon pathways and inflammation response in monocytes (Figure 8C & 8D). Mass cytometry analysis of whole blood from MPN patients revealed the inhibition of MAPK signaling pathway in monocytes following α-KG treatment (Figure 8E). Taken together, these findings suggest that α-KG inhibition of monocyte activation and inflammation might also contribute to its therapeutic effects in MPNs.

## Discussion

Thrombosis and bleeding complications are common in MPNs, but the underlying pathophysiology involved in these events is incompletely understood. Some reports have suggested platelet hyperreactivity and enhanced granule secretion as a potential mechanism for thrombosis in the setting of MPN (14, 19, 35–39). In this study, we used a multiomic profiling approach to characterize MPN platelets and uncovered functional, transcriptional and metabolic signatures associated with platelet hyperreactivity. Importantly, we identified the PI3K/AKT/mTOR signaling pathway as a significant contributor to the platelet dysregulation observed in MPN patients. We further demonstrated that both direct inhibition of the PI3K/AKT/mTOR signaling pathway and metabolic intervention via α-KG supplementation suppress platelet activation. Importantly, our results suggest therapeutic potential of α-KG supplementation to prevent platelet hyperreactivity in MPN patients.

Consistent with our results, a recent large-scale platelet RNA-seq profiling also suggested altered immune, metabolic, and proteostatic pathways in all three MPN subtypes, with the MPN platelet transcriptome robustly predicting disease progression, therefore highlighting the importance of platelet biology in MPNs (40). Although PI3K/AKT/mTOR signaling has been established as a key regulator of proliferation, cancer, longevity and mitochondrial homeostasis, its role in mediating platelet hyperreactivity in MPNs has not been previously reported (41). Our multiomic and functional results identified PI3K/AKT/mTOR signaling as a key driver of platelet hyperreactivity in MPNs. The effects of dual PI3K/mTOR inhibition by omipalisib underscores the essential role of PI3K/AKT/mTOR signaling in mediating platelet metabolism and hyperactivation in MPNs, suggesting potential therapeutic role in MPNs (42, 43). We also identified metabolic alterations of enhanced OXPHOS and glycolysis activity in MPN platelets. We further demonstrate that α-KG supplementation is an effective metabolic intervention for MPNs, inhibiting both PI3K/AKT/mTOR signaling and mitochondrial activation. Thus, our work reveals PI3K/AKT/mTOR signaling and metabolic alterations as the major drivers of platelet reactivity in MPNs.

Metabolic alterations in MPNs have been previously reported. It has been shown that glutaminase inhibitors suppresses the growth of *JAK2 V617F* mutant cell lines and MPN patient CD34+ cells (44). IDH2 inhibitors showed efficacy in cells from MPN patients carrying both *JAK2* and *IDH2* mutations (45). Recently, elevated glycolysis in *JAK2*-mutant HSPCs was identified as a novel target to treat MPNs (46). Our multiomic results suggest enhanced OXPHOS activities in MPN platelets, validated by increased rates of basal respiration and ATP generation observed in bioenergetic analysis. In this work, we showed that mitochondrial abnormalities fuel platelet hyperactivation in MPNs with both increased basal respiration and reserve capacity after TRAP6 stimulation. Of note, our results suggest that the PI3K/AKT/mTOR signaling serves as a driver of mitochondrial abnormalities in MPN platelets; as the inhibition of mitochondrial activities by α-KG supplementation decreased pathway activation.

α-KG has been shown to be highly versatile in regulating cellular activities, as evidenced by the extensive list of described α-KG-dependent proteins, including Jumonji domain-containing histone demethylases, TET proteins mediating DNA methylation and Prolyl-hydroxylase domain enzymes degrading HIF proteins (26). Our results show inhibition of PI3K/AKT/mTOR signaling by α-KG supplementation, however we cannot exclude potential alternative mechanisms. Further research is needed to explore targets of α-KG in human cells. Lastly, α-KG has been previously reported to maintain embryonic stem cell pluripotency via epigenetic regulation, but its effects on hematopoiesis have been unexplored (47). In this study we demonstrated inhibition of myeloid differentiation in *Jak2 V617F* transplanted mice but not in wild type mice, indicating the effectiveness and safety of α-KG supplementation in the MPN setting. We also observed inhibition of monocyte activation and cytokine secretion by α-KG treatment, in agreement with the previously described role of α-KG in alleviating inflammation and oxidative stress (48, 49). Therefore, the inhibition of myeloid differentiation, which resulted in reduced platelet, RBC and monocyte counts, in α-KG-treated mice could also be a result of decreased chronic inflammation in MPNs.

In summary, our data reveal a previously unrecognized mitochondrial disorder in platelets from MPN patients; this energetic alteration leads to platelet hyperreactivity potentially contributing to thrombotic events (Figure 8F). We also identified aberrant PI3K/AKT/mTOR signaling partially rectified by α-KG supplementation (Figure 8F). These findings may lead to novel therapeutic approaches targeting platelet hyperreactivity and chronic inflammation in MPN.

## Methods

### Cell culture

MEG-01 (ATCC) and UKE-1 (Coriell Institute) cells were cultured in RPMI 1640 Medium ATCC modification and RPMI 1640 Medium (ThermoFisher), respectively, supplemented with 10% fetal bovine serum (FBS) and 1% penicillin/streptomycin. All cell lines were maintained at 37°C and 5% CO2 and regularly tested for mycoplasma.

### Blood collection and platelet isolation from human

Whole blood was drawn into 4.5 mL tubes containing buffered sodium citrate in accordance with an Institutional Review Board-approved protocol at Washington University in St. Louis. Blood was transferred into 15ml tubes with the addition of 10% (v/v) prewarmed citrate-dextrose solution (ACD). Platelet-rich plasma (PRP) was separated from the other cellular components of blood by centrifugation for at 250g for 20 min without brake and then carefully withdrawn using a plastic Pasteur pipette without disturbing buffy coat. Prostaglandin E1 (PGE1) was added to PRP to a final concentration of 1 μM to prevent aggregation and then spun again for 10 min at 1000 g without brake. The platelet pellet was then carefully washed with 5 mL of Tyrode’s-ACD (9 parts of Modified Tyrode’s buffer and 1 part of ACD in the presence of PGE1) and spun for 5 min at 700 g. The resultant pellet was gently resuspended in Modified Tyrode’s buffer (129 mM NaCl, 0.34 mM Na2HPO4, 2.9 mM KCl, 12 mM NaHCO3, 20 mM HEPES, 5 mM glucose, 1 mM MgCl2; pH 7.3) and adjusted to a count of 3 x 10^8^ platelets/ml for following experiments.

### PLA and platelet activation analysis by flow cytometry

PLA were tested as previously described (50). Whole blood samples were diluted 5-fold by adding HEPES buffer (145 mM NaCl, 5 mM KCl, 1 mM MgSO4, 0.5 mM NaH2PO4, 5 mM glucose, 10 mM HEPES/Na) within 30 minutes of collection. Diluted blood samples were incubated with PerCP-conjugated CD61 and Pacific Blue-conjugated anti-CD45 mAb for PLA analysis. Samples were also stained with APC-conjugated anti-CD62P mAb for platelet CD62P expression analysis. Samples were stained for 10 min at room temperature. For platelet activation, 1 μM TRAP6 or 5 μg/ml collagen stimulation was added simultaneously with antibody staining. Next, samples were fixed with 1.5 % paraformaldehyde, followed by dilution of 4.6-fold with distilled water for red cell lysis. Samples were analyzed by flow cytometry.

### Platelet aggregation assay

Light transmission aggregometry of washed human platelets (300 x 10^3^/μl) was performed in PAP-8E platelet aggregometer (Biodata corporation). Aggregations were induced with TRAP6 or collagen as indicated in each experiment.

### scRNA-seq and analysis

All MPN patients in the cohort were recruited at the Hematology Department of Washington University School of Medicine in St. Louis. Sample processing were performed as previously published (51, 52). Chromium Next GEM Single cell 3′ Reagent v3.1 (Dual Index) was used for GEM Generation, barcoding and cDNA library preparation per manufacturer’s guidelines, and cDNA was sequenced using the 10X Genomics Single-cell RNA-Seq platform at the Genome Technology Access Center at the McDonnell Genome Institute of Washington University in St. Louis. scRNA-seq analysis details are provided in Supplemental Methods.

### Sample preparation for metabolomics and proteomics analysis

Washed platelets were isolated as described above without adding glucose or other nutrients in Modified Tyrode’s buffer. 3 x 10^8^ platelets were carefully washed with PBS for twice without flushing or pipetting the pellet to remove all out-sourced metabolites. Carefully remove the supernatant and flash freeze the pellet with liquid nitrogen. Metabolomics and proteomics were performed as previously published and details are provided in Supplemental Methods (53).

### Seahorse assay

Bioenergetics of washed platelets (20 × 10^6^/well) were determined by Seahorse XFe96 (Agilent Technologies) as previously described (54). After measurement of basal OCR, OCR due to proton leak was determined by Oligomycin A (2.5 μM) treatment. Maximal uncoupled OCR was measured by the addition of the uncoupler carbonyl cyanide p-(trifluoro-methoxy) phenyl-hydrazone (FCCP; 0.5 μM). Nonmitochondrial OCR (defined as the OCR of all cellular processes excluding mitochondrial respiration) was measured in the presence of Rotenone/antimycin A (1 μM). In subsets of samples, TRAP6 (20 μM) was added before Oligomycin A to measure TRAP6-stimulated energy demand. In subsets of samples, platelets were pre-incubation with Octyl-α-KG (200 μM) or other drugs as indicated for one hour to determine their effects on platelet bioenergetics.

### Western Blot

Washed platelets were lysed with RIPA buffer (Thermo Fisher Scientific) with the existence of protease and phosphatase inhibitor cocktail (Thermo Fisher Scientific) and quantified using Bradford assay (Thermo Fisher Scientific). 20 μg boiled protein were loaded for detection as previously published (55). Quantification of western blot bands were performed using imageJ software.

### Platelet adhesion and spreading assay

Platelets were isolated from *Jak2* V617F knock-in mice by sequential centrifuges. Platelets were incubated with 250 μM Octyl-α-KG at 37°C or DMSO control for one hour followed by 100 μM PAR-4 agonist peptide for 5 minutes. Treated platelets were incubated on fibrinogen-coated coverslips for 45 minutes at 37°C in wells of 24-well plate. Coverslips were washed three times with PBS, fixed with paraformaldehyde, permeabilized with 0.1% Triton X-100 and mounted with VECTASHIELD antifade mounting medium with phalloidin (Vector laboratories) and imaged using Nikon A1Rsi confocal microscope and NIS-Elements AR software. Number and area of attached platelets on the coverslips were quantified with CellProfiler software (n = 9).

### Colony forming unit (CFU) assays

CFU assays were performed in semi-solid culture using Methocult M3434 (StemCell Technologies, Vancouver, BC, Canada), containing IL-3, IL-6, SCF and EPO. Sorted cKIT+ cells were seeded at 1000 cells/ml with 250 μM Octyl-α-KG or DMSO control in triplicate. Colonies were counted 14 days post-seeding.

### Megakaryocyte differentiation and proplatelet formation

CD34+ cells were sorted from MPN patients and normal individuals cryopreserved BMMCs with MicroBead (Miltenyi Biotec). 2 x 10^4^ sorted cells were plated in triplicate in serum-free media containing StemSpan™ Megakaryocyte Expansion Supplement (STEMCELL Technologies) for 10-12 days. Media were replaced every 5 days. Megakaryocyte cell surface markers, including FITC-conjugated CD41 and PerCP-conjugated CD61, were measured by flow cytometry. MEG-01 cells were treated with 10 ng/mL phorbol 12-myristate 13-acetate (PMA) to induce megakaryocyte differentiation (56).

### Plasma cytokine analysis

Peripheral blood plasma collected under standard protocols was stored at −80 °C. Concentrations of 30 cytokines/chemokines were analyzed in duplicate using the Meso Scale Discovery platform with the V-PLEX human cytokine 30-plex kit (Meso Scale Discovery). Statistical analysis was performed using GraphPad Prism (GraphPad Software).

### In vivo models

#### Wild type mice

Regular water or 1% dietary α-KG was administered to 7 week-old C57BL/6 CD45.2 mice. Mice were treated for four weeks and peripheral blood was collected every week for hemavet analysis.

#### *Jak2* V617F model

cKIT+ cells from CD45.2 *Jak2 V617F* donor mice were transplanted into CD45.1 lethally irradiated recipient mice as previously published and details are provided in Supplemental Methods (52, 57). Two weeks after transplant, Mice were supplemented with regular water or 1% dietary α-KG for 6 weeks. Peripheral blood was collected every other week for hemavet tests and. Mice were sacrificed at endpoint, and body, spleen, and liver weights were recorded. Bone marrow samples were collected for flow cytometry and bulk RNA-seq analysis. Femur bones were collected for hematoxylin and eosin staining.

For platelet activation experiments, *Jak2 V617F* mice were supplemented with regular water (n = 7) or 2% α-KG (n = 7) for one week. Peripheral blood was collected for platelet activation analysis as described above.

### Statistics

Statistical analyses were performed using GraphPad Prism and R software. Two-tailed Student’s t test, Mann Whitney U test, two-way ANOVA and Pearson correlations were performed as indicated. All relevant assays were performed independently at least 3 times.

### Study approval

Patient and healthy individual control peripheral blood (PB) or bone marrow (BM) samples were obtained with written consent according to a protocol approved by the Washington University Human Studies Committee (WU no. 01-1014) and to the Helsinki Declaration of the World Medical Association. Mononuclear cells (PBMC or BMMC) were obtained by Ficoll gradient extraction and cryopreserved according to standard procedures. Additional BMMCs were purchased from STEMCELL Technologies (Vancouver, Canada). Lists of patient samples utilized in this study were provided in Table 1 and Table 3-6. All mice procedures were conducted in accordance with the Institutional Animal Care and Use Committee of Washington University (no. 20-0463).

## Data availability

Bulk RNA-seq, scRNA-seq data, and mass cytometry data will be made publicly available upon publication. More detailed information for this paper can be found in the supplementary materials. Additional data information is available upon reasonable request from the authors.

## Acknowledgements

This work was supported by NIH grants R01HL134952 (S.T.O.). Technical support was provided by the Alvin J. Siteman Cancer Center Tissue Procurement Core Facility, Flow Cytometry Core, and Immunomonitoring Laboratory, which are supported by NCI Cancer Center Support Grant P30CA91842. The Immunomonitoring Laboratory is also supported by the Andrew M and Jane M Bursky Center for Human Immunology and Immunotherapy Programs. We thank D. Bender, R. Lin, and K. Link for assistance with mass cytometry experiments. Bulk RNA-seq and scRNA-seq were performed at Genome Technology Access Center core at the McDonnell Genome Institute (GTAC@MGI). We thank Mary Fulbright for assistance with mouse colony management.

## Author contributions

F.H., A.B.A.L., T.K., A.L. and L.Y. performed experiments. K.J.A., N,M.L., D.A.C.F., L.A.H. and M.C.F. provided technical support. M.J.C. coordinated clinical sample collection. A.D.A. performed proteomics and metabolomics experiments. F.H., S.M.S., J.D.P., and S.T.O. designed and supervised the experiments. F.H., J.D.P., and S.T.O. wrote the manuscript. All authors read and approved of the manuscript.

## Supplementary methods

### Bulk RNA-seq analysis

RNA from mouse bone marrow and human purified CD14+ monocytes were extracted using RNeasy Mini Kit (Qiagen) with DNase treatment in duplicate. Samples were then indexed, pooled, and sequenced by Illumina NovaSeq 6000. Basecalls and demultiplexing were performed with Illumina’s bcl2fastq software and a custom python demultiplexing program with a maximum of one mismatch in the indexing read. RNA-seq reads were then aligned to the Ensembl release 76 primary assembly with STAR (2.5.1a). Genes expressed in less than 30 percent of samples were excluded from further analysis. TMM normalization size factors were calculated with EdgeR package (3.36.0) to adjust for samples for differences in library size. The TMM size factors and the matrix of counts were then use for differential expression analysis between conditions and the results were filtered for only those genes with adjusted p-values less than or equal to 0.05. GSEA was performed using software from Broad Institute (http://software.broadinstitute.org/gsea/index.jsp) as described previously (18). Published expression data were downloaded from Gene Expression Omnibus (https://www.ncbi.nlm.nih.gov/geo/query/acc.cgi?acc=gse2006).

### scRNA-seq analysis

scRNA-Seq data were initially processed by the Cell Ranger pipeline (v2.1.0) from 10 X Genomics using the human reference genome (GrCH38). The generated count matrices were then analyzed with the Seurat (4.3.0) package in R (4.1.3). Read counts were normalized to library size, scaled by 10,000, log transformed, and filtered based on the following criteria: cells with less than 700 genes detected, proportion of the UMIs mapped to mitochondrial genes over 0.2 were excluded from analysis; Cell-cell variation in gene expression driven by the number of detected molecules and mitochondrial gene expression were regressed out using linear regression. Dimensionality reduction, PCA, UMAP projection, and graph-based clustering analysis were then performed. Marker genes for each cluster were identified with Wilcoxon rank sum test. Gene set enrichment analysis of DEGs was run using fgsea and the Hallmark gene set from msigdbr. Gene sets for cell scoring were downloaded from MSIGDB (www.gsea-msigdb.org) (58, 59).

### Metabolomics

Metabolites were extracted from frozen platelet pellets at 4 x 10^8^/mL by vigorous vortexing in the presence of ice cold 5:3:2 MeOH:MeCN:water (v/v/v) for 30 min at 4°C. Plasma samples were thawed on ice then a 20 μL aliquot was treated with 480 μL of the aforementioned solution then vortexed for 30 min at 4°C. Supernatants were clarified by centrifugation (10 min, 12,000 g, 4°C). The resulting extracts were analyzed (10 μL per injection for platelets, 20 μL per injection for plasma) by ultra-high-pressure liquid chromatography coupled to mass spectrometry (UHPLC-MS - Vanquish and Q Exactive, Thermo). Metabolites were resolved on a Kinetex C18 column (2.1 x 150 mm, 1.7 μm) using a 5-minute gradient method exactly as previously described (60). Following data acquisition, .raw files were converted to .mzXML using RawConverter then metabolites assigned and peaks integrated using Maven (Princeton University) in conjunction with the KEGG database and an in-house standard library. Quality control was assessed as using technical replicates run at beginning, end, and middle of each sequence as previously described. Data were adjusted by protein quantity from proteomics. For data analysis, the peak intensities were normalized by median, log transformed and Pareto scaled on MetaboAnalyst website (www.metaboanalyst.ca). Processed data were analyzed with R using limma package.

### Proteomics

The samples were processed using the S-Trap filter (Protifi) according to the manufacturer’s procedure. Briefly, a pellet after metabolomics extraction was solubilized in 50 μl of 5% SDS. Samples were reduced with 10 mM DTT at 55°C for 30 min, cooled to room temperature, and then alkylated with 25 mM iodoacetamide in the dark for 30 min. Next, a final concentration of 1.2% phosphoric acid and then six volumes of binding buffer (90% methanol; 100 mM triethylammonium bicarbonate, TEAB; pH 7.1) were added to each sample. After gentle mixing, the protein solution was loaded to a S-Trap filter, spun at 1,000 x g for 1 min, and the flow-through collected and reloaded onto the filter. This step was repeated three times, and then the filter was washed with 200 μL of binding buffer 3 times. Finally, 1 μg of sequencing-grade trypsin (Promega) and 150 μL of digestion buffer (50 mM TEAB) were added onto the filter and digested carried out at 37°C for 6 h. To elute peptides, three stepwise buffers were applied, with 100 μL of each with one more repeat, including 50 mM TEAB, 0.2% formic acid in H2O, and 50% acetonitrile and 0.2% formic acid in H2O. The peptide solutions were pooled, lyophilized and resuspended in 100 μl of 0.1% FA.

A 20 μl of each sample was loaded onto individual Evotips for desalting and then washed with 20 μL 0.1% FA followed by the addition of 100 μL storage solvent (0.1% FA) to keep the Evotips wet until analysis. The Evosep One system (Evosep) was used to separate peptides on a Pepsep column, (150 μm inter diameter, 15 cm) packed with ReproSil C18 1.9 μm, 120A resin. The system was coupled to the timsTOF Pro mass spectrometer (Bruker Daltonics) via the nano-electrospray ion source (Captive Spray, Bruker Daltonics).

The mass spectrometer was operated in PASEF mode. The ramp time was set to 100 ms and 10 PASEF MS/MS scans per topN acquisition cycle were acquired. MS and MS/MS spectra were recorded from *m/z* 100 to 1700. The ion mobility was scanned from 0.7 to 1.50 Vs/cm2. Precursors for data-dependent acquisition were isolated within ± 1 Th and fragmented with an ion mobility-dependent collision energy, which was linearly increased from 20 to 59 eV in positive mode. Low-abundance precursor ions with an intensity above a threshold of 500 counts but below a target value of 20000 counts were repeatedly scheduled and otherwise dynamically excluded for 0.4 min.

MS/MS spectra were extracted from raw data files and converted into .mgf files using MS Convert (ProteoWizard, Ver. 3.0). Peptide spectral matching was performed with Mascot (Ver. 2.6) against the Uniprot human database. Mass tolerances were +/- 15 ppm for parent ions, and +/- 0.4 Da for fragment ions. Trypsin specificity was used, allowing for 1 missed cleavage. Met oxidation, protein N-terminal acetylation and peptide N-terminal pyroglutamic acid formation were set as variable modifications with Cys carbamidomethylation set as a fixed modification.

Scaffold (version 5.0, Proteome Software) was used to validate MS/MS based peptide and protein identifications. Peptide identifications were accepted if they could be established at greater than 95.0% probability as specified by the Peptide Prophet algorithm. Protein identifications were accepted if they could be established at greater than 99.0% probability and contained at least two identified unique peptides. Data were analyzed with R using DEP package (1.16.0). Normalized data were input for gene set enrichment analysis.

#### Mass cytometry (CyTOF)

Whole blood was drawn into 4.5 mL tubes containing sodium heparin in accordance with an Institutional Review Board-approved protocol at Washington University in St. Louis. 200 μl whole blood were incubated with 200 μM Octyl-α-KG or DMSO for 1 hour followed by the stimulation of 10 ng/ml TNFα for 5 min. Add 1 x Proteomic Stabilizer from Smart Tube Inc (1.4ml of Stabilizer per ml of blood), mix gently and incubate for 10 minutes at room temperature. Keep samples at -80°C until it is ready to be stained. Frozen samples were thawn in cold water bath (10°C to 15°C) with agitation of the water for approximately 20 minutes until the samples are fully thawed. Sampels were washed with Thaw-Lyse Buffer to lyse erythrocytes accordingly. Cells were then permeabilized, barcoded, stained with surface marker antibodies, permeabilized with methanol, stained with intracellular antibodies, and resuspended in DNA IR-intercalator. CyTOF experiments were conducted on a CyTOF2 mass cytometry (Fluidigm) with validated antibody panels. Post CyTOF run, cell identities were debarcoded and data was analyzed in Cytobank (cytobank.org). Monocytes were gated as follows: CD45+, CD66b-, CD14+. Raw median signal intensities were used for analysis.

