## supplemental figures for "Multiomic Profiling Reveals Metabolic Alterations Mediating Aberrant Platelet Activity and Inflammation in Myeloproliferative Neoplasms"

### Figure S1

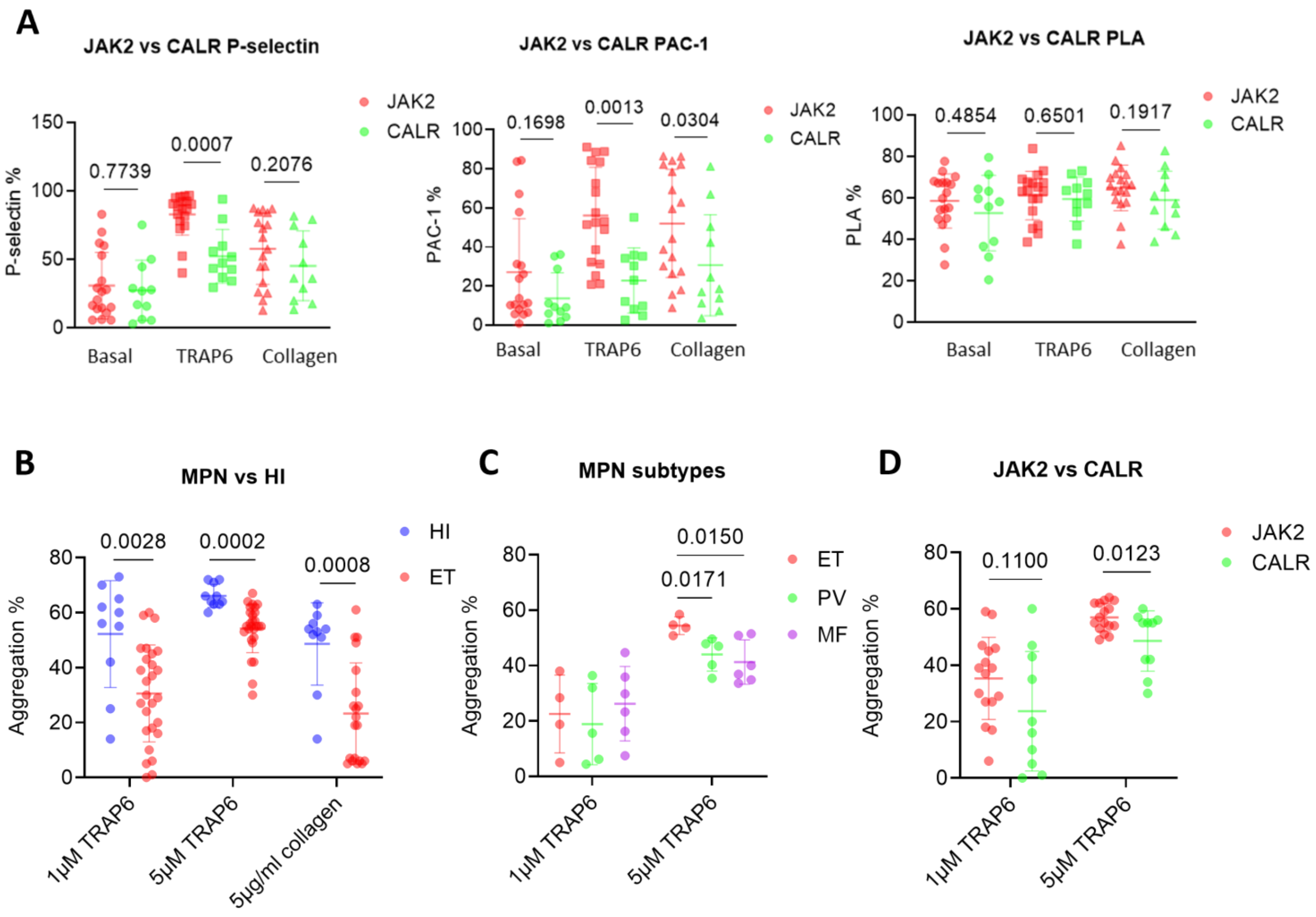


**Figure S1. Analysis of platelet activation, secretion and aggregation.**

A) Dot plot comparing P-selectin, αIIbβ3 integrin expression and PLA formation of platelets from peripheral blood of ET patients with JAK2 and CALR mutations by flow cytometry.

B) Dot plot showing maximal aggregation intensity of washed platelets from HIs and MPN patients. Data is shown as mean ± SD.

C) Dot plot showing maximal aggregation intensity of washed platelets from subjects with different MPN subtypes.

D) Dot plot showing maximal aggregation intensity of washed platelets from MPN patients with JAK2 and CALR mutations.

### Figure S2

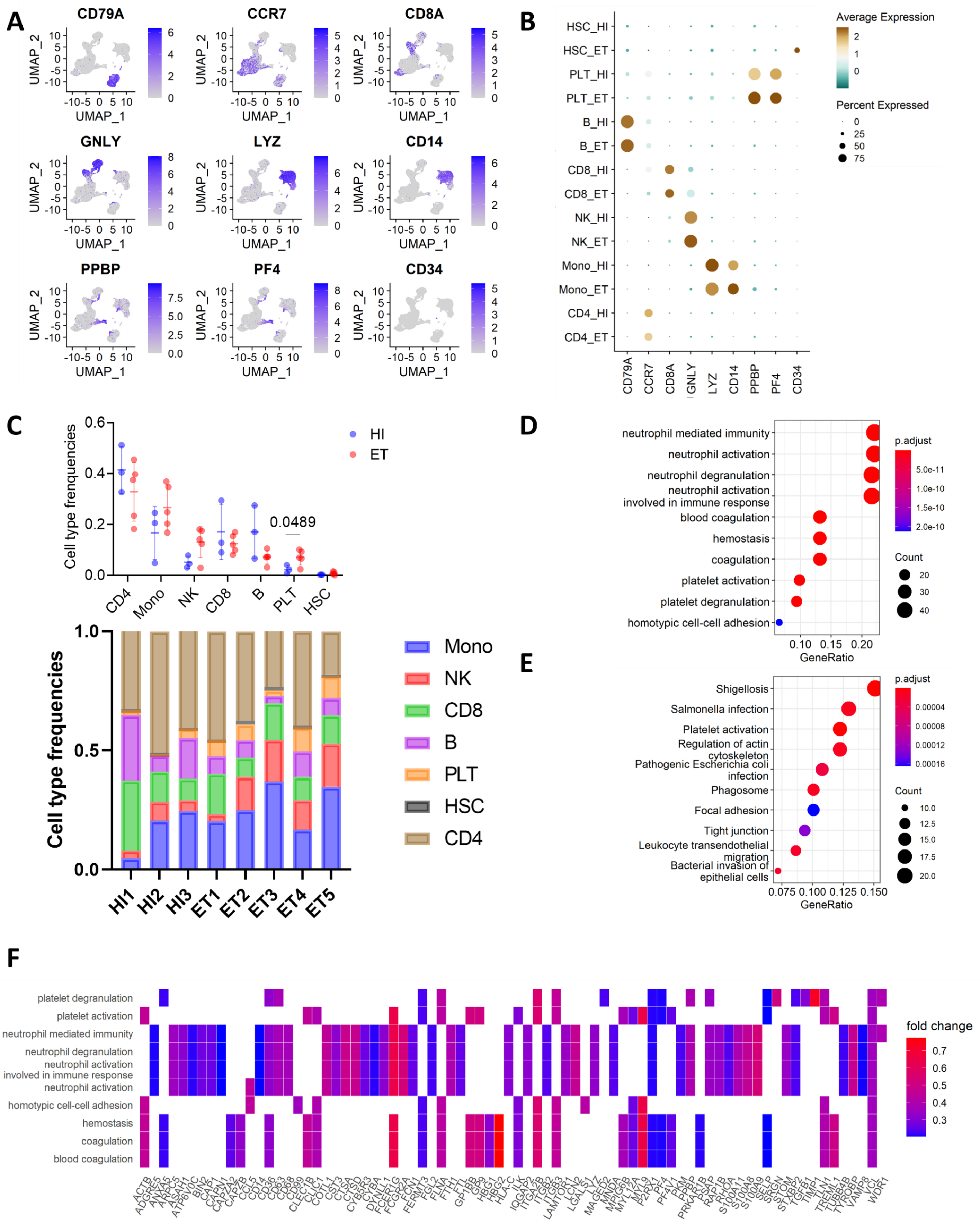

A
CD79A
CCR7
CD8A
GNLY
LYZ
CD14
PPBP
PF4
CD34
UMAP_1
UMAP_2
B
HSC_HI
HSC_ET
PLT_HI
PLT_ET
B_HI
B_ET
CD8_HI
CD8_ET
NK_HI
NK_ET
Mono_HI
Mono_ET
CD4_HI
CD4_ET
Average Expression
Percent Expressed
C
Cell type frenquencies
HI
ET
0.0489
CD4
Mono
NK
CD8
B
PLT
HSC
Cell type frequencies
Mono
NK
CD8
B
PLT
HSC
CD4
HI1
HI2
HI3
ET1
ET2
ET3
ET4
ET5
D
neutrophil mediated immunity
neutrophil activation
neutrophil degranulation
neutrophil activation involved in immune response
blood coagulation
hemostasis
coagulation
platelet activation
platelet degranulation
homotypic cell-cell adhesion
GeneRatio
p.adjust
Count
E
Shigellosis
Salmonella infection
Platelet activation
Regulation of actin cytoskeleton
Pathogenic Escherichia coli infection
Phagosome
Focal adhesion
Tight junction
Leukocyte transendothelial migration
Bacterial invasion of epithelial cells
GeneRatio
p.adjust
Count
F
platelet degranulation
platelet activation
neutrophil mediated immunity
neutrophil degranulation
neutrophil activation involved in immune response
neutrophil activation
homotypic cell-cell adhesion
hemostasis
coagulation
blood coagulation
fold change

### Figure S2

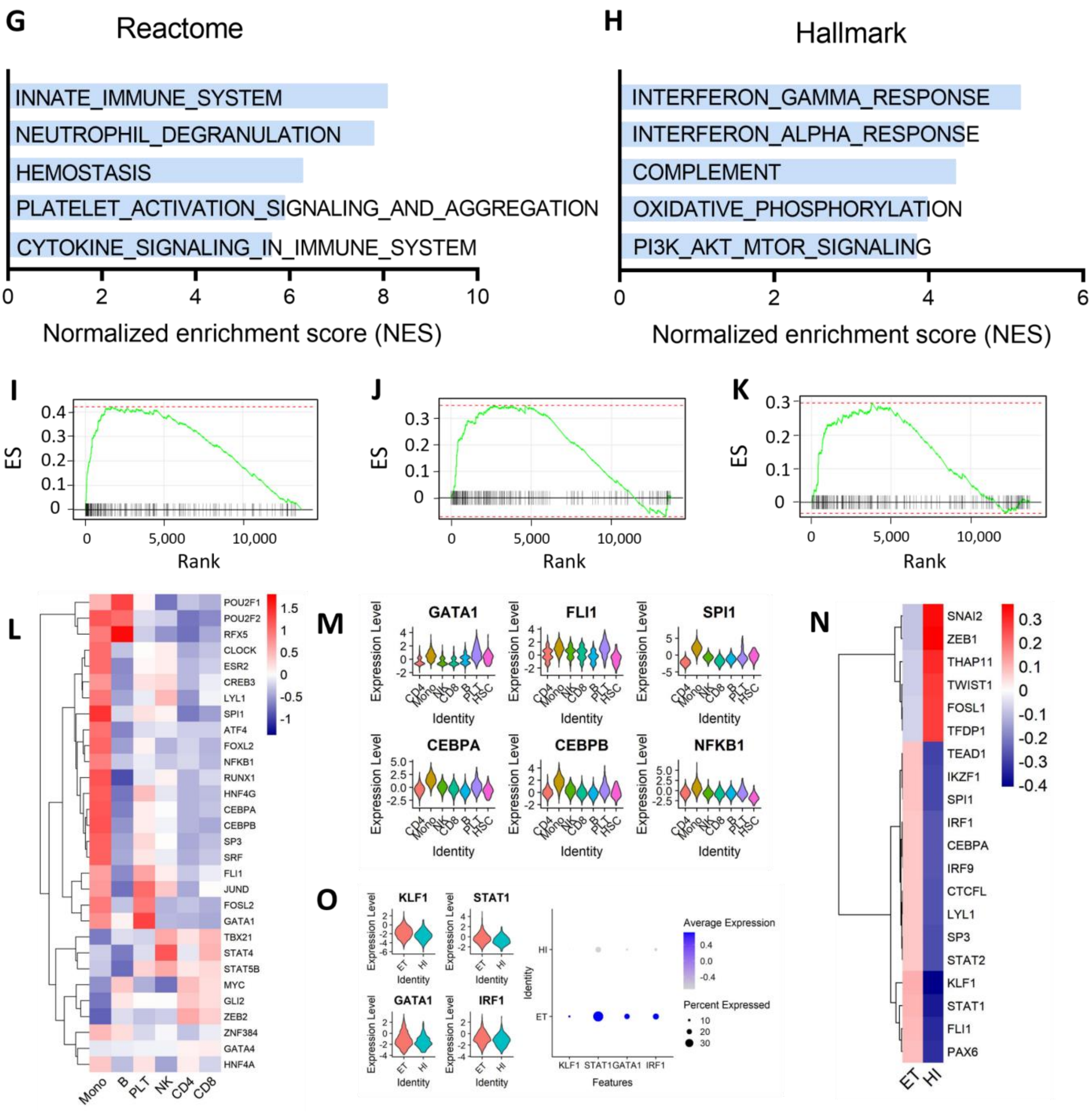
G
Reactome
INNATE_IMMUNE_SYSTEM
NEUTROPHIL_DEGRANULATION
HEMOSTASIS
PLATELET_ACTIVATION_SIGNALING_AND_AGGREGATION
CYTOKINE_SIGNALING_IN_IMMUNE_SYSTEM
0 2 4 6 8 10
Normalized enrichment score (NES)
H
Hallmark
INTERFERON_GAMMA_RESPONSE
INTERFERON_ALPHA_RESPONSE
COMPLEMENT
OXIDATIVE_PHOSPHORYLATION
PI3K_AKT_MTOR_SIGNALING
0 2 4 6
Normalized enrichment score (NES)
I
J
K
ES
Rank
0 5,000 10,000
L
POU2F1
POU2F2
RFX5
CLOCK
ESR2
CREB3
LYL1
SPI1
ATF4
FOXL2
NFKB1
RUNX1
HNF4G
CEBPA
CEBPB
SP3
SRF
FLI1
JUND
FOSL2
GATA1
TBX21
STAT4
STAT5B
MYC
GLI2
ZEB2
ZNF384
GATA4
HNF4A
Mono B PLT NK CD4 CD8
1.5 1 0.5 0 -0.5 -1
M
GATA1
FLI1
SPI1
CEBPA
CEBPB
NFKB1
Expression Level
Identity
CD4 Mono NK CD8 B PLT HSC
O
KLF1
STAT1
GATA1
IRF1
ET HI
Average Expression
Percent Expressed
10 20 30
Features
N
SNAI2
ZEB1
THAP11
TWIST1
FOSL1
TFDP1
TEAD1
IKZF1
SPI1
IRF1
CEBPA
IRF9
CTCFL
LYL1
SP3
STAT2
KLF1
STAT1
FLI1
PAX6
ET HI
0.3 0.2 0.1 0 -0.1 -0.2 -0.3 -0.4

**Figure S2. scRNA-seq analysis.**

A) Feature plots of genes for cell type identification.

B) Dot plot of top differentially expressed genes for cell type identification.

C) Dot plot and stacked bar plot showing frequency of differernt type of cells in all 8 samples for scRNA-seq.

D) GO enrichment analysis of differentially-expressed genes (DEGs) in platelets from ET patients.

E) KEGG enrichment analysis of DEGs in platelets from ET patients.

F) Heatmap showing the distribution of DEGs in GO enrichment pathways.

G) Barplot showing top 5 reactome pathways enriched in platelets from ET patients.

H) Barplot showing top 5 hallmark pathways enriched in platelets from ET patients.

I) GSEA enrichment plots for "reactome platelet activation signaling and aggregation" gene set enriched in platelets from ET patients.

J) GSEA enrichment plots for "hallmark IFN-γ response" gene set enriched in platelets from ET patients.

K) GSEA enrichment plots for "hallmark OXPHOS" gene set enriched in platelets from ET patients.

L) Heatmap showing top 20 most variable transcriptional factors (TFs) in each type of cells based on VIPER scores on DoRothEA's regulons.

M) Violin plots showing the expression of representative TFs in each type of cells.

N) Heatmap showing top 20 most variable TFs in platelets from ET compared to platelets from HI.

O) Violin plots and dot plots showing the expression of representative TFs in platelets from ET and HI.

### Figure S3

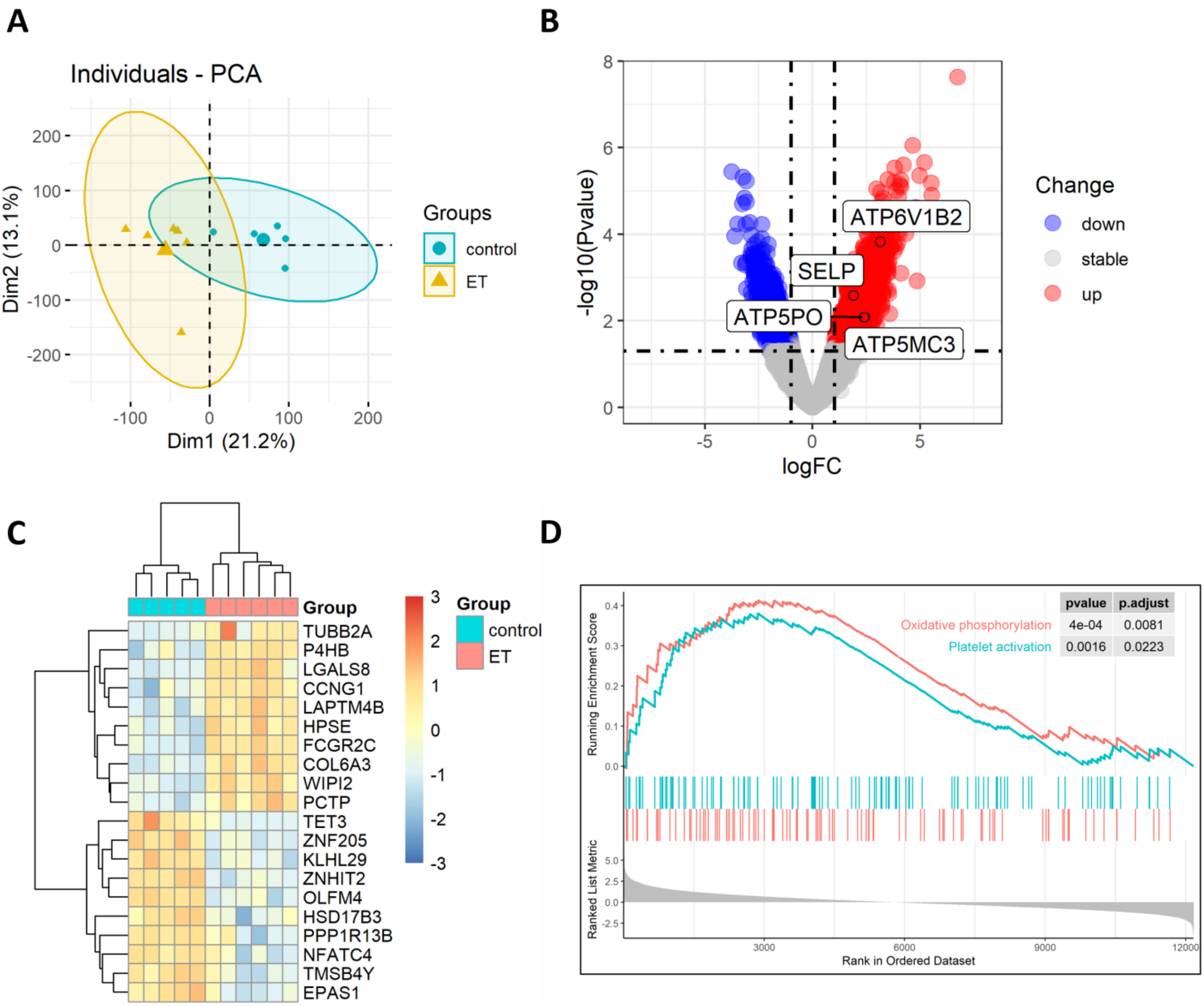


**Figure S3. Analysis of GSE2006 microarray between platelets from HIs and ET patients.**

A) PCA plot of platelets from ET (n = 6) and HIs (n = 5) in GSE2006.

B) Volcano plot showing differentially expressed genes (DEG) between HIs and ET patients in platelets in GSE2006.

C) Heatmap of top 10 DEGs in HIs and ET patients in GSE2006.

D) GSEA enrichment plots for “hallmark OXPHOS” and “reactome platelet activation signaling and aggregation” gene sets enriched in platelets from ET patients.

### Figure S4

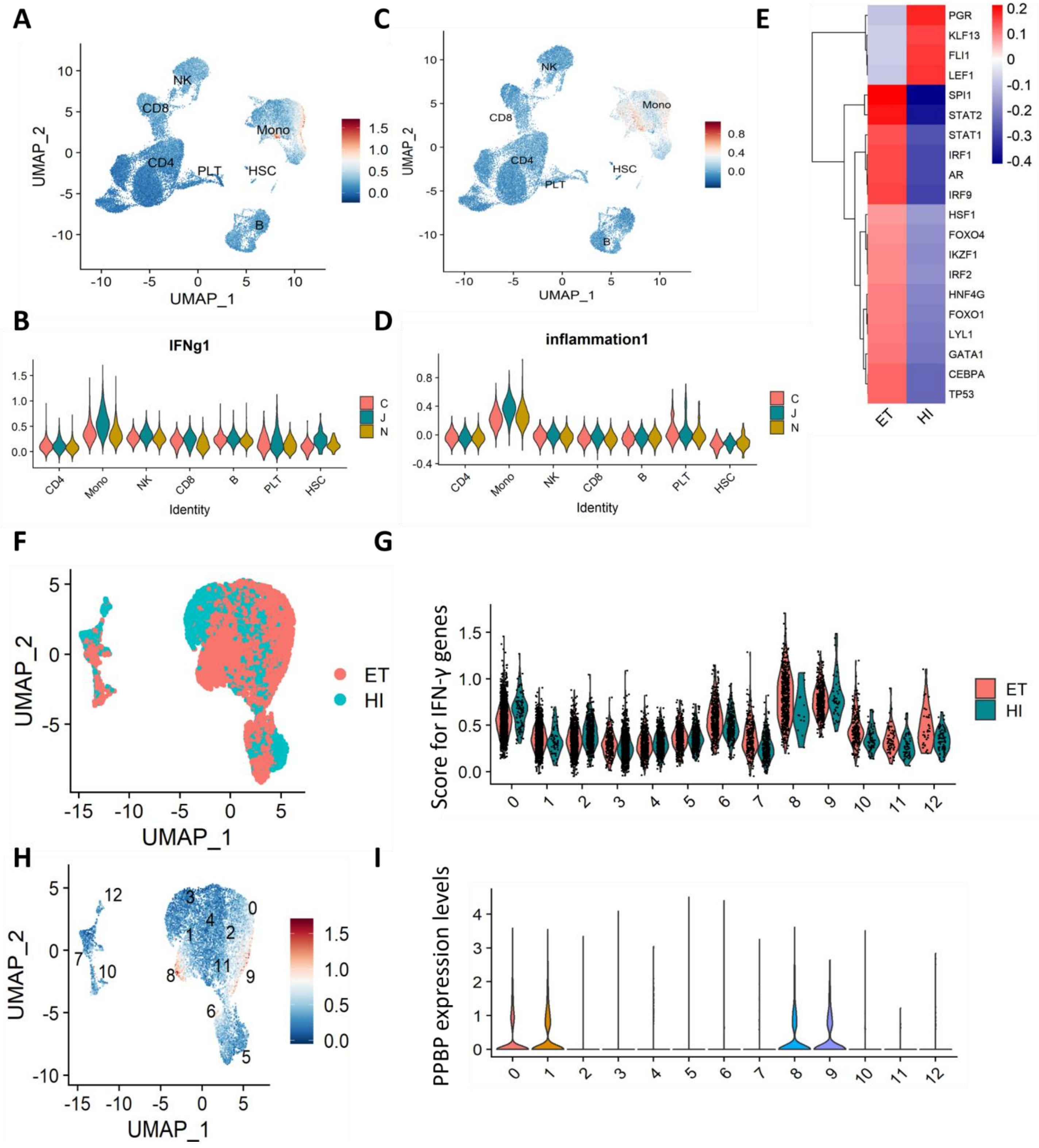
A
B
C
D
E
F
G
H
I
UMAP_1
UMAP_2
NK
CD8
Mono
CD4
PLT
HSC
B
IFNg1
inflammation1
Identity
C
J
N
PGR
KLF13
FLI1
LEF1
SPI1
STAT2
STAT1
IRF1
AR
IRF9
HSF1
FOXO4
IKZF1
IRF2
HNF4G
FOXO1
LYL1
GATA1
CEBPA
TP53
ET
HI
Score for IFN-γ genes
PPBP expression levels

**Figure S4. scRNA-seq analysis of monocytes.**

A) UMAP plot showing scores for genes in “hallmark interferon gamma response” gene set.

B) Violin plot showing scores for genes in “hallmark interferon gamma response” gene set (C: ET patients with CALR mutations; J: ET patients with JAK2 mutation; N: HIs).

C) UMAP plot showing scores for genes in “hallmark inflammatory response” gene set.

D) Violin plot showing scores for genes in “hallmark inflammatory response” gene set (C: ET patients with CALR mutations; J: ET patients with JAK2 mutation; N: HIs).

E) Heatmap showing top 20 most variable TFs in monocytes from ET compared to monocytes from HI.

F) UMAP plot showing scores for genes in “hallmark interferon gamma response” gene set in monocytes from HIs and ET patients.

G) Violin plot showing scores for genes in “hallmark interferon gamma response” gene set in monocytes from HIs and ET patients.

H) UMAP plot showing PPBP (a marker for platelets) expression levels in monocytes from HIs and ET patients.

I) Violin plots showing PPBP (a marker for platelets) expression levels in monocytes from HIs and ET patients.

### Figure S5

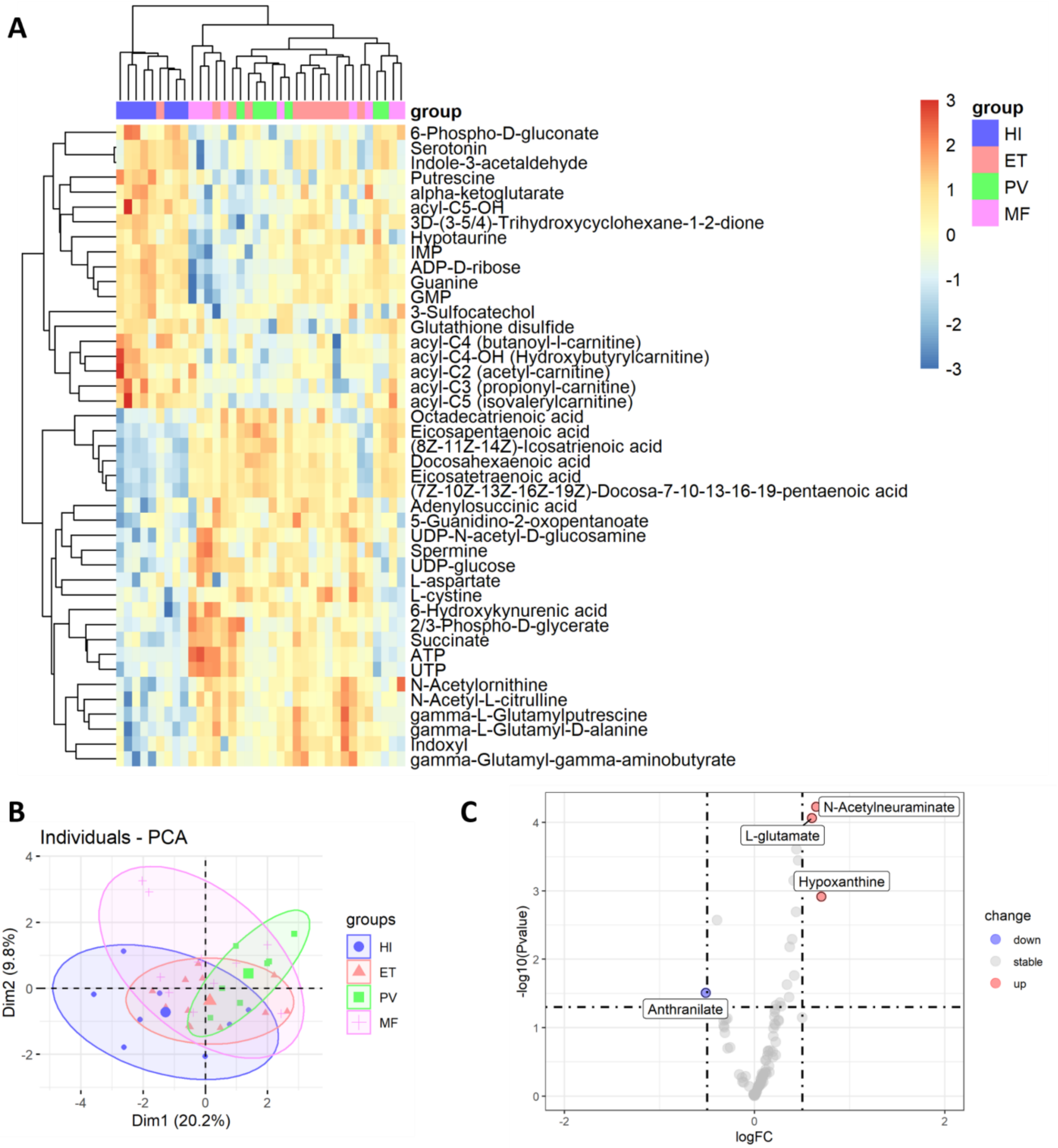


**Figure S5. Plasma metabolite LC-MS showed no difference between His and MPNs.**

A) Heatmap of down- (n = 19) and up-regulated (n = 24) metabolites in platelets between ET patients and HIs (P<0.05, |fold change| > 2^0.5).

B) PCA plot of metabolites in plasma from MPN patients and HIs.

C) Volcano plot of plasma DAMs between MPN patients and HIs.

### Figure S6

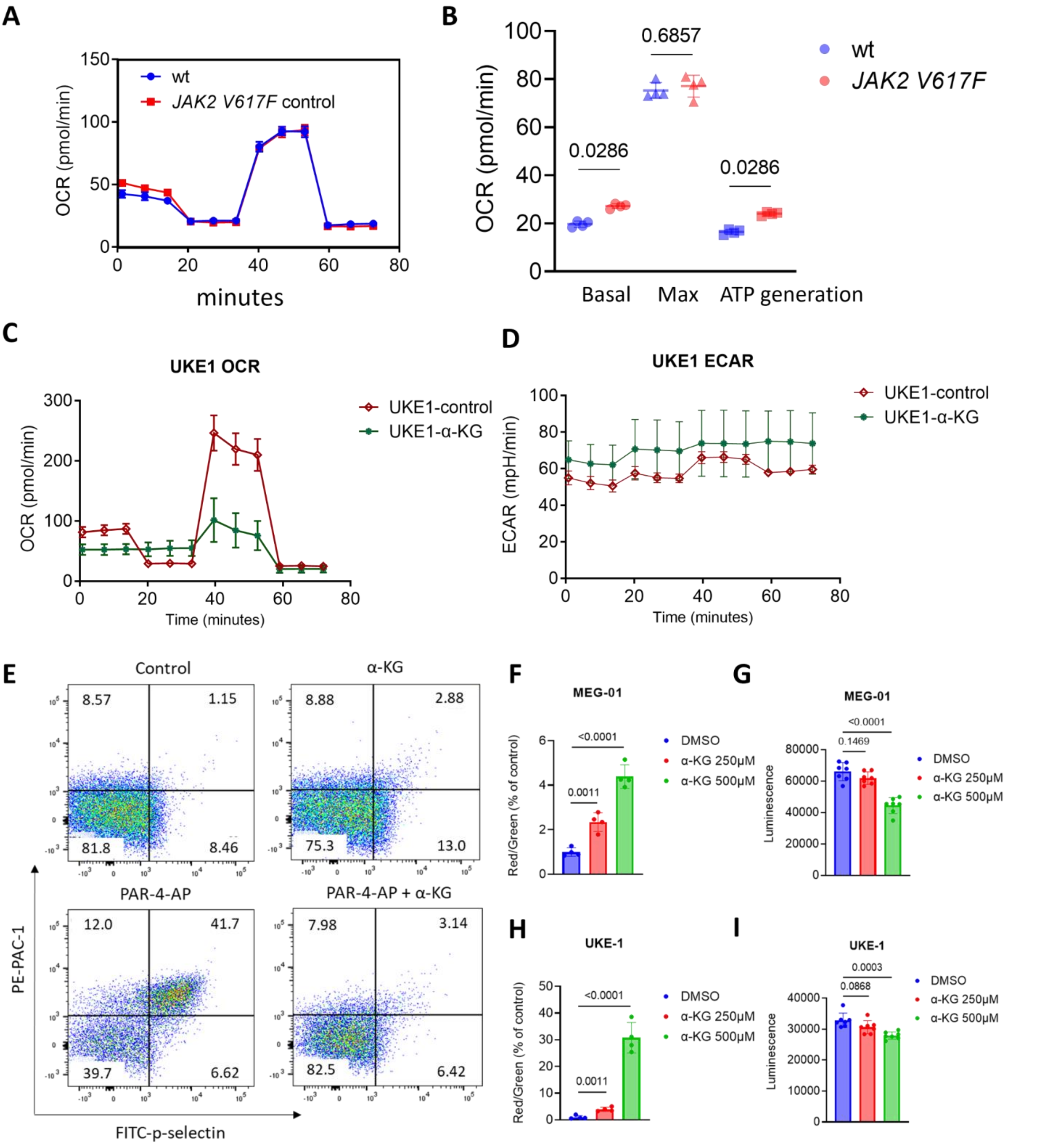

A
OCR (pmol/min)
wt
JAK2 V617F control
minutes
B
OCR (pmol/min)
0.0286
0.6857
0.0286
wt
JAK2 V617F
Basal
Max
ATP generation
C
UKE1 OCR
OCR (pmol/min)
UKE1-control
UKE1-α-KG
Time (minutes)
D
UKE1 ECAR
ECAR (mpH/min)
UKE1-control
UKE1-α-KG
Time (minutes)
E
Control
8.57
1.15
81.8
8.46
α-KG
8.88
2.88
75.3
13.0
PAR-4-AP
12.0
41.7
39.7
6.62
PAR-4-AP + α-KG
7.98
3.14
82.5
6.42
PE-PAC-1
FITC-p-selectin
F
MEG-01
Red/Green (% of control)
<0.0001
0.0011
DMSO
α-KG 250μM
α-KG 500μM
G
MEG-01
Luminescence
<0.0001
0.1469
DMSO
α-KG 250μM
α-KG 500μM
H
UKE-1
Red/Green (% of control)
<0.0001
0.0011
DMSO
α-KG 250μM
α-KG 500μM
I
UKE-1
Luminescence
0.0003
0.0868
DMSO
α-KG 250μM
α-KG 500μM

**Figure S6. Effects of α-KG on mouse platelets and megakaryocytic cell lines.**

A) Representative OCR profiles of platelets from wild type and *JAK2 V617F* knock-in mice.

B) Quantification of mitochondrial respiration parameters (n = 4) of platelets from wild type and *JAK2 V617F* knock-in mice. Data are mean ± SD. Statistics were assessed by two-tailed Mann-Whitney U test.

C) Representative OCR profiles of UKE-1 cells with the incubation of Octyl-α-KG or DMSO.

D) Representative ECAR profiles of UKE-1 cells with the incubation of Octyl-α-KG or DMSO.

E) Representative images showing effects of α-KG on activation of washed platelets from *JAK2 V617F* knock-in mice. Washed platelets from *JAK2 V617F* knock-in mice were treated with Octyl-α-KG for 1 hour followed by PAR-4-AP stimulation and flow cytometry analysis.

F) Bar plot showing the effects of α-KG on mitochondrial membrane potential in MEG-01 cells determined by JC-1 dye staining.

G) Bar plot showing the effects of α-KG on intracellular ATP level in MEG-01 cells determined by ATPlite luminescence assay.

H) Bar plot showing the effects of α-KG on mitochondrial membrane potential in UKE-1 cells determined by JC-1 dye staining.

I) Bar plot showing the effects of α-KG on intracellular ATP level in UKE-1 cells determined by ATPlite luminescence assay.

### Figure S7

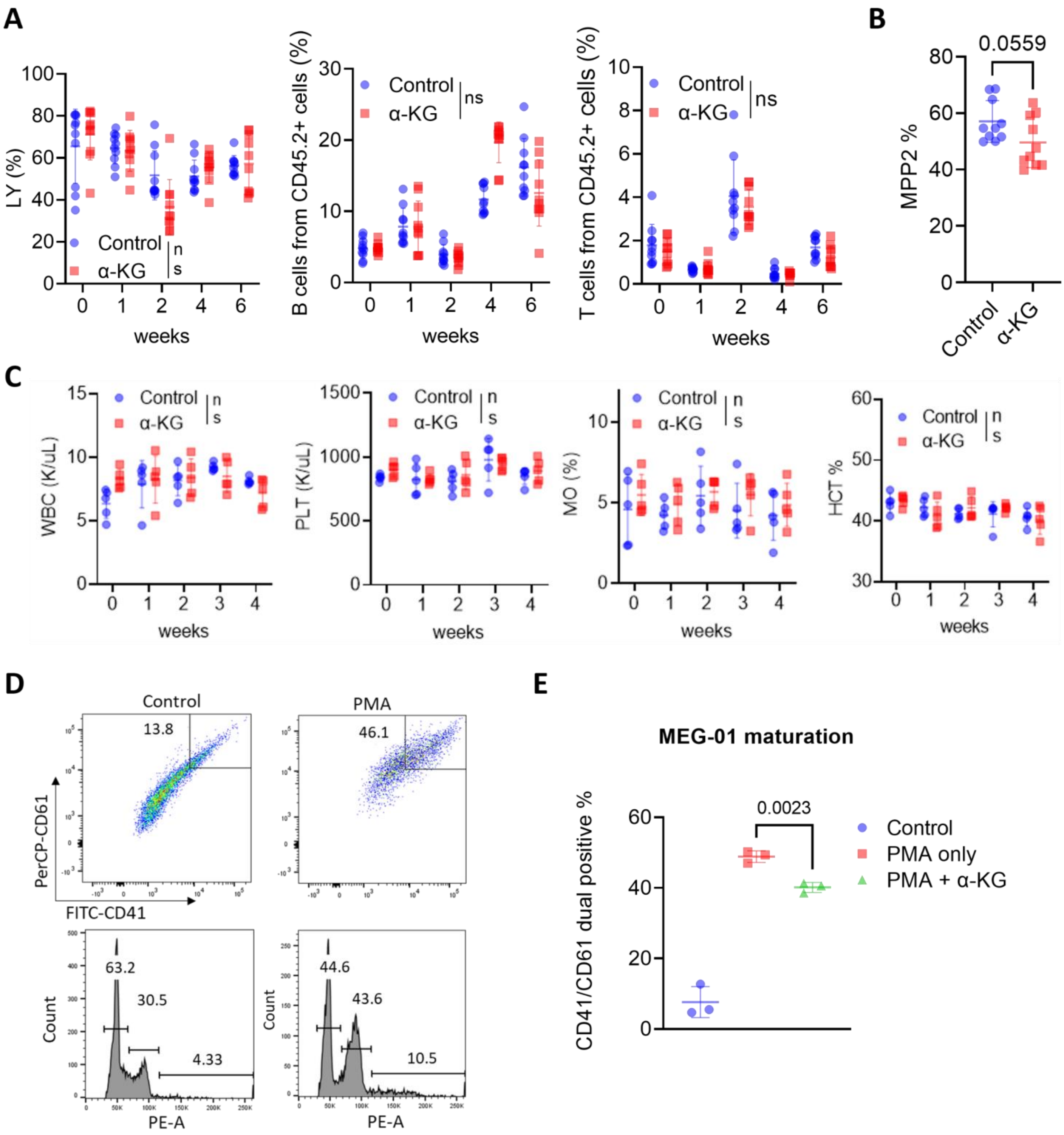
A
LY (%)
Control
α-KG
ns
weeks
B cells from CD45.2+ cells (%)
T cells from CD45.2+ cells (%)
B
0.0559
MPP2 %
Control
α-KG
C
WBC (K/uL)
PLT (K/uL)
MO (%)
HCT %
D
Control
PMA
13.8
46.1
PerCP-CD61
FITC-CD41
63.2
30.5
4.33
44.6
43.6
10.5
Count
PE-A
E
MEG-01 maturation
0.0023
CD41/CD61 dual positive %
Control
PMA only
PMA + α-KG

**Figure S7.** ***in vivo*** **treatment of α-KG inhibited MPN progression.**

A) Lymphocytes (LY), B and T cell ratios from CD45.2+ cells of transplanted mice treated with vehicle or α-KG across multiple timepoints. Data are mean ± SD. Statistics were assessed by two-way ANOVA with Dunnett's multiple comparisons test ( ns: not significant, $P > 0.05$).

B) Percentage of MPP2 cells of transplanted mice treated with vehicle or α-KG across multiple timepoints. Data are mean ± SD. Statistics were assessed by Two-tailed Student's t test.

C) WBC, platelet counts, monocytes and HCT ratios from C57BL/6J mice treated with vehicle (n = 5) or α-KG (n = 5) across multiple timepoints. Data are mean ± SD. Statistics were assessed by two-way ANOVA with Dunnett's multiple comparisons test ( ns: not significant, $P > 0.05$).

D) Representative images showing the maturation of MEG-01 cells with PMA treatment (CD41/CD61 expression and ploidy analysis by flow cytometry)

E) Quantification of percentages of CD41/CD61 dual positive cells with α-KG treatment. Data are mean ± SD. Statistics were assessed by Two-tailed Student's t test. MEG-01 cells were treated with phorbol myristate acetate (PMA) for 2 days with 500 μM Octyl-α-KG or DMSO control. The ratio of CD41- and CD61-dual positive cells were determined by flow cytometry (n =3). Ploidy analysis was determined by propidium iodide (PI) staining.

### Figure S8

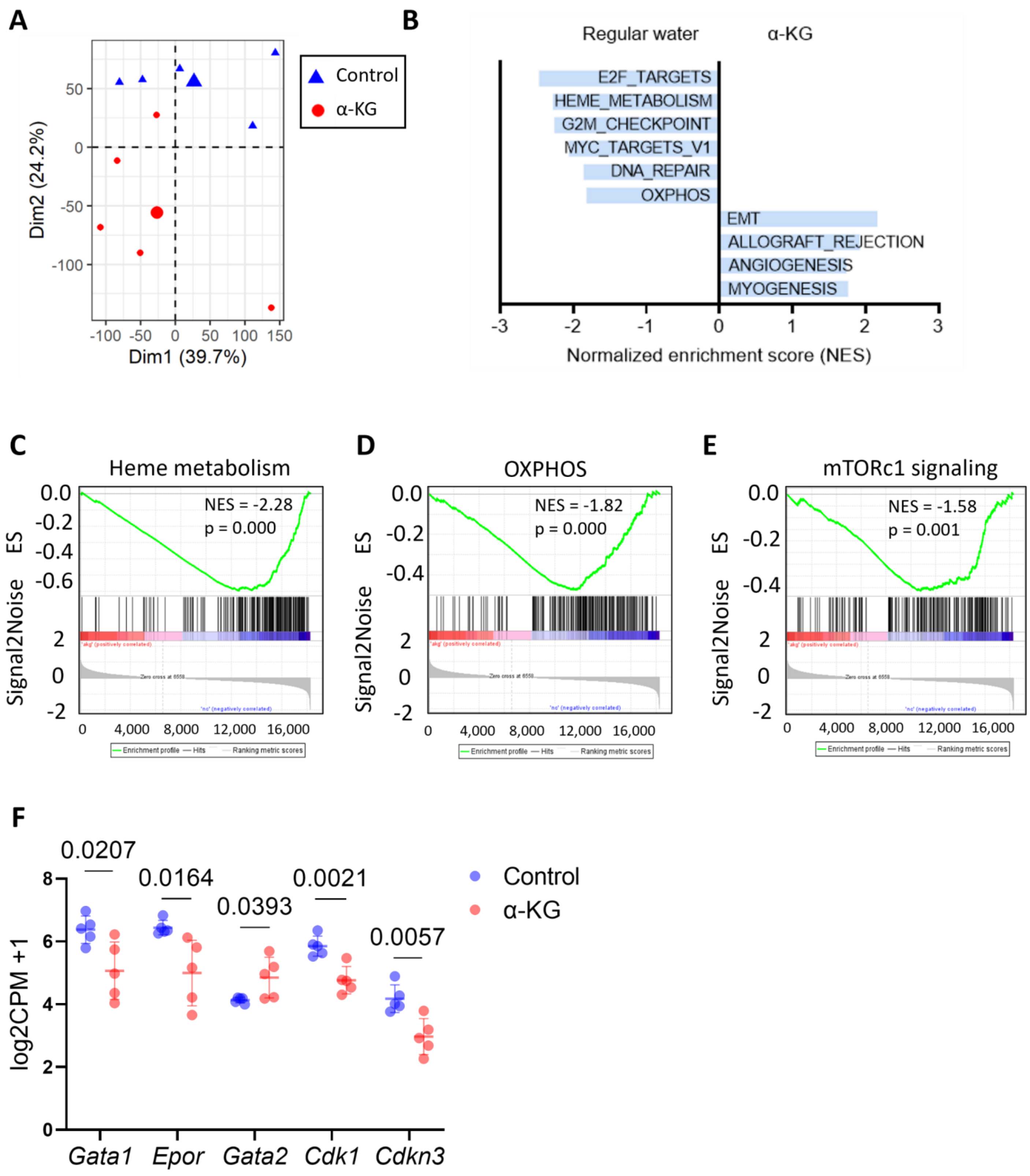
A
Control
α-KG
Dim1 (39.7%)
Dim2 (24.2%)
B
Regular water
α-KG
E2F_TARGETS
HEME_METABOLISM
G2M_CHECKPOINT
MYC_TARGETS_V1
DNA_REPAIR
OXPHOS
EMT
ALLOGRAFT_REJECTION
ANGIOGENESIS
MYOGENESIS
Normalized enrichment score (NES)
C
Heme metabolism
NES = -2.28
p = 0.000
D
OXPHOS
NES = -1.82
p = 0.000
E
mTORc1 signaling
NES = -1.58
p = 0.001
ES
Signal2Noise
Enrichment profile — Hits — Ranking metric scores
F
0.0207
0.0164
0.0393
0.0021
0.0057
Control
α-KG
log2CPM +1
Gata1
Epor
Gata2
Cdk1
Cdkn3

**Figure S8. Analysis of bulk RNA-seq of bone marrow from control (n = 5) and 1% α-KG-supplemented mice (n = 5).**

A) PCA plot for bulk RNA-seq data of bone marrow from control and 1% α-KG-supplemented mice.

B) Barplot of top 10 hallmark pathways with the highest GSEA enrichment scores when comparing between control and 1% α-KG-supplemented groups.

C) GSEA enrichment plots for “Oxidative phosphorylation” gene set enriched in control vs. α-KG-supplemented mice.

D) GSEA enrichment plots for “Heme metabolism” gene set enriched in control vs. α-KG-supplemented mice.

E) GSEA enrichment plots for “mTORc1 signaling” gene set enriched in control vs. α-KG-supplemented mice.

F) Dot plot of RNA-seq counts of representative genes.
